# A Multigenerational “Dirty” Mouse Model for Studying Trauma-Induced Immune Dysregulation and Infection Susceptibility

**DOI:** 10.1101/2025.06.18.660185

**Authors:** Alexandria Byskosh, John Pulford, Ekaterina Murzin, Zoe Michael, Chyna Gray Lovell, Bailin Niu, Anupamaa Seshadri, Gabriel Brat, Ashley Zajdel, Frank Borriello, James Lederer

**Author notes:** Corresponding Author: James Lederer, PhD, 20 Shattuck Street, Thorn Building, Room 425 Boston, MA 02115.

## Abstract

Trauma induces immune dysregulation in both humans and mice, increasing infection susceptibility. Mouse models are critical in research but have been criticized for lacking translational relevance. This study tested whether multigenerational “natural immune” (NI) mice – generated by co-housing C57BL/6 mice with “dirty” pet shop mice and breeding through multiple generations – would develop a more human-like immune response to trauma and infection than “clean” specific pathogen-free (SPF) C57BL/6 mice whose immune systems developed without normal flora. To address this gap, SPF and NI mice underwent burn trauma followed by *Pseudomonas aeruginosa* lung infection. Peripheral blood and bone marrow immune cells were characterized by flow cytometry and mass cytometry (CyTOF). Blood samples from trauma patients were analyzed for comparison. At baseline, NI mice exhibited more neutrophils compared to SPF mice, closer resembling human peripheral immune composition. Following injury, SPF mice demonstrated increased blood neutrophils and monocytes with reduced B and T cells, whereas NI mice exhibited a muted blood immune cell response. In contrast, NI mice showed a robust emergency granulopoiesis response and preserved hematopoietic stem cells (HSCs) following secondary infection, whereas HSCs decreased in SPF mice. Time-matched blood samples from human trauma patients revealed alterations more closely resembling those observed in NI mice. These findings support the hypothesis that NI mice develop a more human-like immune response to trauma and infection than SPF mice. This suggests that NI mice may provide a more translationally relevant platform for studying trauma-induced immune dysfunction and infection susceptibility mechanisms.

**Summary Sentence:** Multigenerational natural immune mice exhibit more human-like immune responses to trauma and infection that offer a model with significant translatable advantages for studying post-injury immune dysfunction in patients.

## Introduction

In the United States, trauma is the leading cause of death among people ages 1-44 and the fourth leading cause of death for people of all ages.^1^ Deaths due to trauma have a tri-modal distribution—*immediate*, *early*, and *late phases*. Infection is the most common complication following trauma and is responsible for *late deaths*, occurring days to weeks following injury.^2–5^ Trauma patients are more susceptible to infection and injury severity is a predominant risk factor.^5,6^ Hospitalized trauma patients who acquire at least one infection have an increased risk of mortality one year following the event.^6,7^

Traumatic injuries create a complex altered host response, resulting in profound immunologic dysfunction and increased risk of infection. During the initial traumatic insult, tissue damage triggers an innate immune response. Following this initial response, innate and adaptive immune systems also activate *pro-* and *counterinflammatory immune responses* with a surge of innate immune cells and dampening of the adaptive immune response with decreased T cell function.^8–11^

At the cellular level, physical insults and systemic infections increase the demand for neutrophils and stimulate a response known as *emergency granulopoiesis—*a process characterized by increased myeloid progenitor cell proliferation and differentiation in the bone marrow.^8,11,12^

Mice are used to model traumatic injury as human and mice share almost 90% of their genetic makeup; however, significant physiologic differences between the two species raise questions regarding the translatability of results.^13^ Current trauma immunology research typically uses specific pathogen-free (SPF) mice which are inbred, genetically uniform, and have standardized microbiomes. In contrast, humans have diverse genetics and microbiomes. Additionally, SPF mice lack natural exposure to common pathogens. Trained immunity is initiated by infection or vaccination, which leads to an enhanced secondary immune response to infection.^14^ The exposure to common pathogens in humans is central to normal immune system development and naturally acquired trained immunity. Therefore, SPF mice, with their altered immune systems, may not be the best model to study human trauma immunology. Studies have shown that SPF mice that were co-housed with pet shop mice can acquire a more human-like immune system, with exposure to “dirty” natural environments influencing immune development and function.^15–17^ Co-housed “dirty” mice have also been shown to exhibit an altered sepsis response, with increased susceptibility to severe polymicrobial infections from cecal ligation and puncture.^14,18^ These mice display heightened Toll-like receptor (TLR)-mediated immune reactions, supporting the concept that co-housed “dirty” mice acquire trained immunity.^14^

To study the immunological consequences of traumatic injury in a potentially more translatable mouse model, we established a multigenerational dirty mouse model platform which we will refer to as natural immune (NI) mice. NI mice are different than co-housed dirty mice as they have undergone natural adaptation to mouse commensal microbiota and pathogens throughout their lifespan and over at least 5 generations, allowing for stable phenotypes.

In this study, we aimed to characterize and compare the immune responses to traumatic injury between SPF and NI mice, with a particular focus on the alterations in peripheral blood and bone marrow immune cells.

Given the differences in immune system exposure and activation between the two groups, we hypothesized that the NI mice would exhibit a more robust and dynamic immune response to injury compared to SPF mice. To test this, we utilized a “two-hit” model of infection to simulate a post-trauma infection scenario. In this model, burn injury was followed by a sublethal *Pseudomonas aeruginosa* lung infection, which is clinically relevant as pneumonia is a common hospital-acquired infection in trauma patients.^3,5,19–21^ Using mass cytometry (CyTOF) and flow cytometry, we compared the blood and bone marrow of SPF and NI mice with respect to immune cell subsets and marker profiles. We found that NI mice displayed distinct baseline and injury-induced immune profiles compared to SPF mice, with more stable immune cell populations following burn injury, altered cytokine responses, and neutrophil and monocyte phenotypes that more closely resemble those seen in human trauma patients—highlighting the potential of NI mice as a more clinically relevant model for studying trauma-induced immune dysregulation and infection susceptibility.

## Materials and Methods

### Mice

Inbred, male C57BL/6 were purchased from Charles River Laboratories (Wilmington, MA, USA) as our “clean” specific pathogen-free (SPF) mice. These mice were maintained in our full-barrier animal facility under controlled temperature, humidity, and 12-hour light-dark regimen. These mice were provided with standard feed and water *ad libitum* and were acclimated for at least two weeks prior to use. Our “dirty” or natural immune (NI) mice were inbred, male C57BL/6 initially purchased through Charles River Laboratories. Female C57BL/6 mice were originally co-housed with different color female mice purchased from a pet shop supplier. After co-housing female mice for 12 weeks, the “dirty” C57BL/6 female mice were mixed with male C57BL/6 mice to breed. The first-generation offspring were then bred to establish an active multigenerational NI C57BL/6 breeding colony. NI mice were bred for up to 5 generations prior to use in this study. Microbial taxa were profiled in SPF and NI mice using the Transnetyx metagenomics platform with One Codex profiling (Cordova, TN, USA) **(Supplementary Figure 1)**. All animal protocols used in this study were approved by the Brigham and Women’s Hospital Standing Committee on Animal Research and found to be in accordance with guidelines set by the US Department of Agriculture and the National Institutes of Health.

### Mouse Burn Trauma Model

Mice were anesthetized with ketamine 125mg/kg and xylazine 20mg/kg via intraperitoneal (i.p.) injection. Once anesthetized, the dorsal fur was shaved, and mice were placed in an insulated plastic device exposing approximately 20% of their total body surface area (TBSA). The exposed region was then submerged in 90°C water bath for 9 seconds, causing a demarcated, full-thickness third degree burn injury. After injury, mice were given 0.05mL of buprenorphine ER at 0.5mg/mL (Wedgewood Pharmacy, Swedesboro, NY, USA) by subcutaneous (s.c.) injection and resuscitated with 1mL saline containing antisedan reversal agent at 0.005mg/ml (Zoetis, Kalamazoo, MI), USA by i.p. injection.

### Mouse Lung Bacterial Infection Model

*P. aeruginosa* bacteria ATCC 27853 (Manassas, VA, USA) were grown with gentle agitation in tryptic soy broth (TSB) medium at 37°C overnight. Frozen stocks were prepared as 110μL aliquots by flash-freezing in liquid nitrogen with long-term storage at -80°C. To prepare bacteria for infection studies, 100μL of bacteria were added to 10mL of TSB and grown for 19 hours at 37°C. Bacteria were harvested by centrifugation at 650 × *g* for 10 minutes and washed once with sterile phosphate buffered saline (PBS). The bacteria were then diluted in sterile PBS to 0.15 absorbance at 600nm wavelength, which reproducibly correlates to 1.4 × 10^9^ CFU/mL. This bacterial suspension was then diluted 1:1 with PBS to achieve a lethal dose of 30% (LD_30_) in uninjured male C57BL/6 SPF mice. For lung infection, mice were anesthetized with ketamine 125mg/kg (Zoetis, Kalamazoo, MI, USA) and xylazine 20mg/kg (Akorn, Lake Forest, IL, USA) by i.p. injection. Mice were then administered a total of 30μL bacterial suspension intranasally. Bacterial CFUs were quantified by drop-plating of serial dilutions on Luria-Bertani (LB) agar plates. Colonies were counted the next morning after incubation at 37°C overnight.

### Mouse Cell Preparations

Cells were prepared from mice three days following burn injury or two days following *P. aeruginosa* infection. Mice were euthanized by CO_2_ asphyxiation. Blood was collected via intracardiac puncture by syringe containing 0.1mL EDTA with PBS and stored in 1mL EDTA tubes to prevent clotting. RBCs were lysed using our own formulated ammonium chloride-based mouse RBC lysis buffer (TL Buffer) that was developed to preserve fragile cells such as neutrophils. Cells were washed twice by centrifugation at 100 × *g* in RPMI1640 containing 5% heat-inactivated fetal bovine serum (FBS), HEPES, non-essential amino acids, L-glutamine, antibiotics/antimycotics, and b-mercaptoethanol (C5 medium). Cell preparations were filtered in 70-μm cell strainers and resuspended in 1mL CryoStore CS10 freezing medium (Biolife Solutions, Bothell, WA, USA). 0.5mL were aliquoted into 2 Cryogenic vials, placed in Mr. Frosty containers for 20 minutes at 4°C, stored at -80° C for a maximum of three months, then transferred to a vapor phase liquid nitrogen (LN_2_) for long term storage.

Bone marrow (BM) was harvested from femurs by removal of the patellar end and placed in a 1.5mL Eppendorf with a hole at the bottom (created with an 18-gauge needle) inserted into 5mL snap-cap tubes. One mL PBS+ 2mM EDTA was added and samples were centrifuged at 1000 × *g* for three minutes. Each femur was then injected with 0.1mL of Liberase TL (0.4mg/mL in PBS, Roche, Indianapolis, IN, USA) and incubated at 37°C for 15 minutes. Digested bones were centrifuged again using the same nested tube setup to collect stromal cells. Pellets were resuspended in 1mL C5 medium, combined with tube rinses, and centrifuged at 100 × *g* for 10 minutes. Red blood cells were lysed in 2mL TL buffer for 5 minutes at room temperature, quenched with 2mL C5, and centrifuged again. Cells were filtered through 70-μm strainers, pelleted at 100 × *g* for 10 minutes, and resuspended in C5 medium for flow cytometry staining.

### Human Cell Preparations

Whole blood was collected in 10mL purple top EDTA tubes from patients three days following traumatic injury and processed within 4 hours of collection. RBCs were lysed using TL Buffer. Cell preparations were centrifuged for 10 minutes at 300 × *g*, resuspended in 10mL C5 medium, and centrifuged for 10 minutes at 200 × *g*. Supernatant was then decanted and cells were resuspended in 2mL Cryostor CS10 freezing medium (Biolife Solutions, Bothell, WA, USA). Aliquots of 0.5mL were placed into Cryogenic vials, and stored as described above. Trauma patient blood samples were collected under informed consent, and all procedures were conducted in accordance with protocols approved by the Institutional Review Board (IRB #2023P002290).

### Flow Cytometry

Freshly isolated mouse BM cells were plated (2.5×10^6^/sample) in 96-well plates. Cells were spun at 750 × g for 3 minutes to pellet for all washes. Wells were stained with 50µL of surface antibody cocktail containing optimal concentrations of individual antibodies and incubated for 30 minutes at room temperature, protected from light. Following surface marker staining, cells were washed twice with 100µL CSB, followed by a single wash with 100µL PBS (ThermoFisher Scientific, Waltham, MA, USA). Cells underwent fixation in 200µL 4% paraformaldehyde (PFA) (Millipore Sigma, Burlington, MA, USA) diluted in PBS for 10 minutes at room temperature. After fixation, cells were resuspended in 200µL PBS and underwent acquisition on a MACSQUANT Analyzer 10 Flow Cytometer (Miltenyi Biotec, Gaithersburg, MD, USA). For data analysis, single color compensation controls were generated for each antibody-fluorophore marker using OneComp eBeads (Thermo Fisher Scientific, Waltham, MA, USA) according to manufacturer’s protocol. Our gating scheme for hematopoietic stem cell subset identifications and flow cytometry antibody panel is provided in **Supplementary Figure 2**.

### Mass Cytometry (CyTOF)

#### Whole Blood CyTOF

Whole blood samples were fixed for 15 minutes in 1.4 volumes of Proteomic Stabilizer (Smart Tube Inc #PROT1-1L) and stored at -80° C. On staining day, samples were thawed at RT and added to 10mL of Thaw/Lyse Buffer (Smart Tube Inc) for 15 minutes at RT to lyse RBCs. Cells were centrifuged at 600 × g for 10 minutes followed by another Thaw/Lyse cycle. Cells were resuspended in 10mL C5 medium, filtered through a 70-µM sieve filter, and centrifuged at 100 × *g.* Cells were plated in 96-well round bottom polypropylene plates at 1-2 × 10^6^ cells/mL per well, pelleted at 750 × g for 3 minutes and permeabilized with the FoxP3/Transcription Factor Staining Buffer Set (Thermo Fisher Scientific, Waltham, MA, USA) for 30 minutes at RT. Following permeabilization, cells were washed twice with PBS and incubated with SCN-EDTA coupled palladium barcoding reagents for 15 minutes at RT followed by 3 washes with PBS and a 15 minute RT incubation with heparin (Sigma-Aldrich, St. Louis, MO, USA).^22^ Cells were then combined, filtered through a polypropylene tube fit with a 40mM filter cap, and centrifuged at 750 × g for 3 minutes. Either Human or Mouse TruStain FcX Fc receptor blocking reagent (BioLegend, San Diego, CA, USA) diluted 1:100 in cell staining buffer (CSB): PBS with 0.5% bovine serum albumin and 0.05 % sodium azide (Sigma Aldrich, St.

Louis, MO, USA) was used to resuspend cells for 10 minutes at RT. The antibody cocktail was added directly to the cells for 1 hour at RT. Mouse and human CyTOF antibody panels are provided in (**Supplementary Table 1A and 1B, respectively**). After incubation with antibodies, cells were washed twice with permeabilization buffer, once with CSB, then fixed for 10 minutes at RT with 4% PFA in PBS. Cells were incubated with an iridium intercalator solution (Standard BioTools, San Francisco, CA, USA) for 20 minutes at RT, then washed and resuspended in Cell Acquisition Solution (CAS+, Standard BioTools, San Francisco, CA, USA). Before acquisition, cells with filtered through a 40-um filter, resuspended at 1 × 10^6^ cells/mL in CAS+, and acquired on a CyTOF XT Mass Cytometer with normalization beads added (Standard BioTools, San Francisco, CA, USA).

#### Total Leukocyte CyTOF

Samples were thawed in a 37°C water bath for 3 minutes and then mixed with 37°C thawing media supplemented with 10U/ml heparin and 25U/mL benzonase nuclease (Sigma Aldrich, St. Louis, MO, USA). 0.5 – 2.0 × 10^6^ cells were stained for 5 minutes with Rhodium viability reagent (Standard BioTools, San Francisco, CA, USA) diluted in culture media. Then 16% stock paraformaldehyde (PFA) (Thermo Fisher Scientific, Hampton, NH, USA) was diluted to 0.8% in PBS and used to fix the cells for 5 minutes. After centrifugation and aspiration, cells were stained with either Human or Mouse TruStain FcX Fc receptor blocking reagent (BioLegend, San Diego, CA, USA) diluted in CSB for 10 minutes followed by addition of cell surface staining antibodies for 30 minutes at RT. All antibodies were prepared and validated by the Harvard Medical Area CyTOF Antibody Resource and Core (Boston, MA, USA). Cells were permeabilized, barcoded, processed, and run on a CyTOF XT Mass Cytometer as described above.

#### CyTOF Data Pre-Processing and Analysis

Acquired samples were bead-normalized on the instrument then the normalized FCS files were gated to remove debris and non-cellular events using OMIQ software (https://www.omiq.ai/). Cleaned data was compensated with a panel specific spillover matrix generated using our 48-marker antibody cocktail on the CyTOF XT instrument. This compensation spillover matrix was generated to subtract cross-contaminating signals from metals using the compensation method in CATALYST in R.^23^ The compensated files were then deconvoluted into individual sample files per panel using a debarcoding algorithm in CATALYST.^23^ All subsequent analyses were run in OMIQ using a workflow for unbiased clustering and cell subset identification using FlowSOM and hierarchical clustering on median marker expression values. UMAP was used for CyTOF staining data visualization in two-dimensions. Statistics for cell and cluster abundance changes were calculated in GraphPad Prism software (GraphPad, San Diego, CA, USA).

### Plasma Cytokine Analysis

Whole blood samples were centrifuged for 20 minutes at 400 × *g* to separate blood cells and plasma. Plasma was collected and stored at -80°C until use. For Luminex bead-based cytokine assays, cytokine standards were diluted 1:2 in Luminex assay incubation buffer (PBS + 0.05% sodium azide + 0.5% bovine serum albumin + 0.05% Triton X-100). 40mL of undiluted plasma was added to a round bottom polystyrene 96-well plate. 40mL of antibody-coupled Luminex bead cocktails were added to each well and plates were incubated at RT with orbital mixing at rate of 300revs/minute. After 1 hour, plates were placed on a magnetic plate to wash twice with Luminex wash buffer (PBS +0.05 sodium azide + 0.05% Triton X-100). Next, 40mL of cytokine detection cocktail diluted in Luminex incubation buffer was added to each well and plates were incubated at RT with orbital mixing at rate of 300revs/minute. After 1 hour, plates were placed on a magnetic plate to wash twice with Luminex wash buffer (PBS +0.05 sodium azide + 0.05% Triton X-100). Next, 40mL of Streptavidin-PE (BioLegend, San Diego, CA, USA) was added to each well and plates were incubated at RT with orbital mixing at rate of 300revs/minute. After 30 minutes, plates were placed on a magnetic plate to wash twice with Luminex wash buffer (PBS +0.05 sodium azide + 0.05% Triton X-100). Each well was suspended in 100mL of Luminex wash buffer and placed in a MAGPIX instrument to generate mean fluorescence intensity (MFI) for cytokine detection on each specific bead region. Our multiplex panel detects the following cytokines: IL-1α, IL-1β, IL-2, IL-4, IL-5, IL-6, IL-10, IL-12p40, IL-12p70, IL-13, IL-17A, IL-18, IL-23, IL-33, IFNγ, TNFα, G-CSF, GM-CSF, M-CSF, SCF, CCL3, CCL4, CCL5, CCL9/10, CXCL3, CXCL5, CXCL7, FLT3L, KC, MIP2, and MCP-1.

### Statistics

GraphPad Prism 10.3.1 software (GraphPad, San Diego, CA, USA) was used for statistical calculations. Two-way ANOVA with multiple comparisons were used to analyze these data, as indicated in the figure legends. For all data, p < 0.05 was considered statistically significant with a 95% confidence interval.

## Results

### Baseline differences in peripheral immune and bone marrow cells between specific pathogen-free (SPF) and natural immune (NI) mice

To evaluate baseline bone marrow (BM) cell populations, detailed analysis of BM hematopoietic stem and progenitor cell (HSPC) populations by flow cytometry was performed using marker expression profiles (**Figure 1A**). To compare baseline peripheral blood immune cell profiles between SPF and NI C57BL/6 mice, we used mass cytometry (CyTOF) as an approach for systems immunology analysis. NI mice had significantly higher levels of circulating neutrophils and monocytes as well as lower levels of B cells than SPF mice (**Figure 1B**).

**Figure 1.**
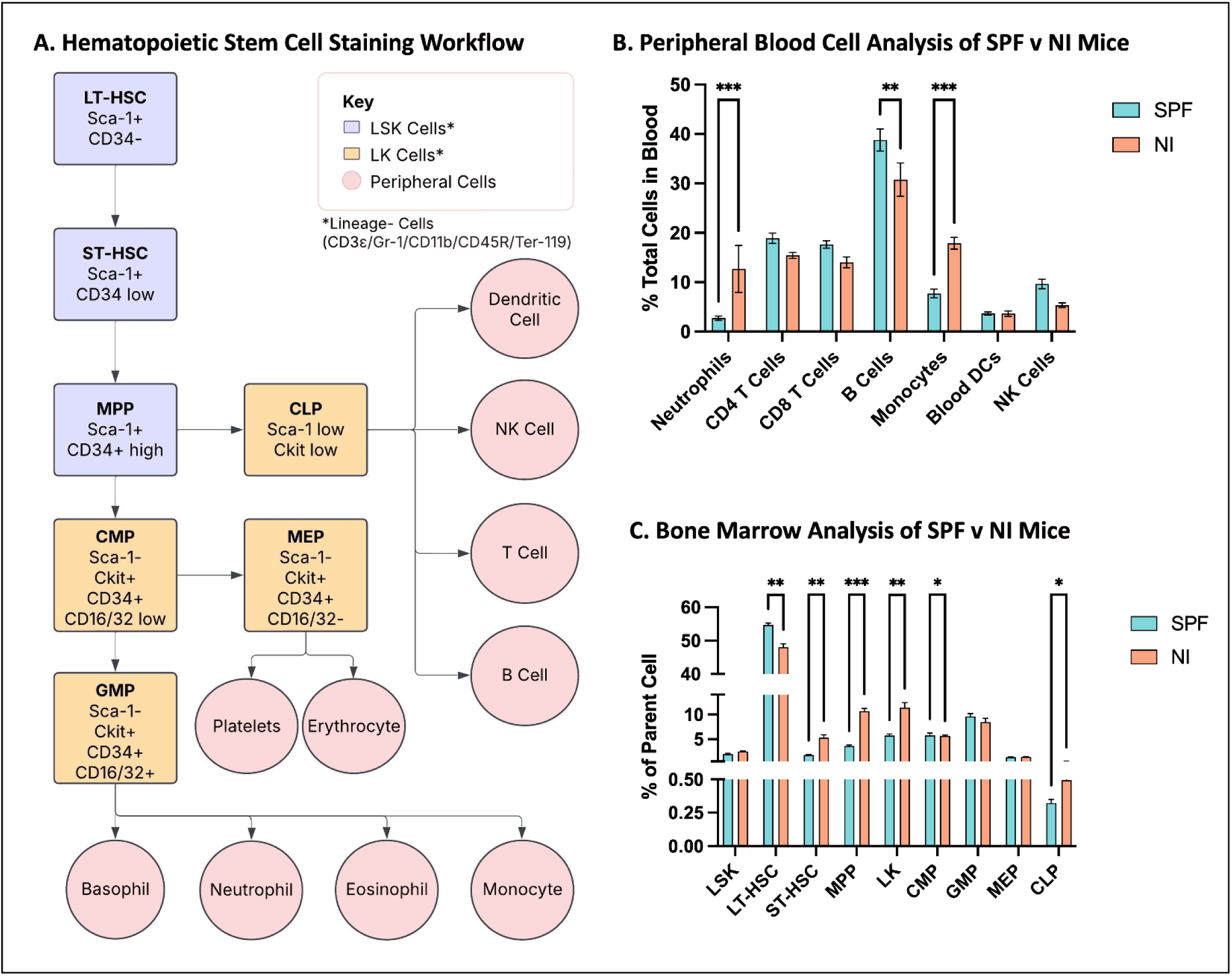
CyTOF and flow cytometry analysis to compare abundances of blood and bone marrow cell populations in Specific Pathogen-Free (SPF) and Natural Immune (NI) mice. (A) Flow chart describing identification of hematopoietic stem cells by flow cytometry staining. (B) Total leukocytes were stained with a 48-marker CyTOF antibody panel to detect baseline relative levels of the indicated immune cell subsets in uninjured C57BL/6 SPF and NI mice. (B) Bone marrow cells were stained by flow cytometry to detect hematopoietic stem and progenitor cells in uninjured C57BL/6 SPF and NI mice. These CyTOF and flow cytometry staining results are plotted as mean +/- SEM and are representative of 3 independent studies using 5 mice per group. Statistical analysis was assessed by two-way ANOVA with multiple comparisons, *= p<0.05, **= p<0.005, ***= p<0.0005, ****=p<0.0001. LSK=Lineage(-)Sca-1(+)c-Kit(-) cells; LT-HSC=Long-Term Repopulating Hematopoietic Stem Cells; ST-HSC=Short-Term Repopulating Hematopoietic Stem Cells; MPP=Multi-Potent Progenitor Cells; LK=Lineage(-)c-Kit(+)Sca-1(-); CMP=Common Myeloid Progenitor Cells; GMP=Granulocyte-Monocyte Progenitor Cells; MEP=Megakaryocyte-Erythrocyte Progenitor Cells; CLP=Common Lymphoid Progenitor Cells.

NI mice showed significantly higher levels of several progenitor cell populations than SPF mice (ST-HSC, MPP, LK, and CLP) except for LT-HSCs, which were significantly lower (**Figure 1C**).

### Characterizing and comparing immune system responses to burn trauma in SPF and NI mice

To evaluate differences or similarities in response to traumatic injury between SPF and NI mice, we used our well-established burn trauma model.^24^ This is a nonlethal traumatic injury that is known to mimic many of the changes seen in human trauma patients. We believe this approach is a robust model to study trauma as it produces a reproducible, survivable injury with longitudinal immune system changes. At 3 days after sham or burn injury, burn-injured SPF mice demonstrated a marked and significant increase in the percentage of blood neutrophils and a decrease in B cells (**Figure 2A-C**). In contrast, NI mice did not show significant changes in neutrophil or B cell populations following burn injury (**Figure 2D**). Burn-injured SPF mice demonstrated a significant increase in circulating monocytes and a decrease in CD4^+^ T cells (**Figure 2C**), while NI mice showed no significant changes in these blood immune cell subsets (**Figure 2D**).

**Figure 2.**
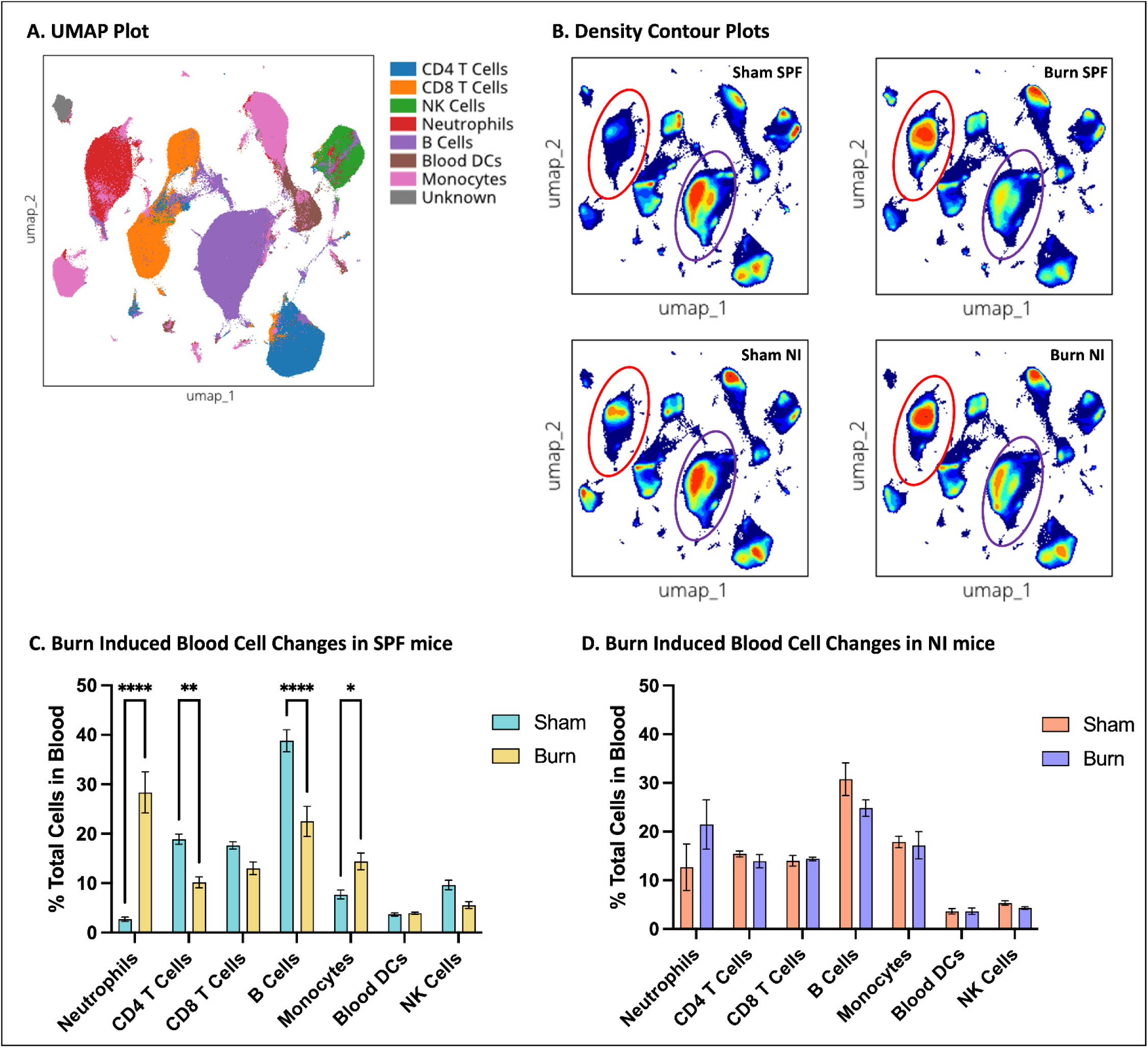
CyTOF analysis to compare abundances of blood cell populations with burn injury in Specific Pathogen-Free (SPF) and Natural Immune (NI) mice. (A) Overlay UMAP plot showing immune cell subset clusters. (B) Density contour plots illustrating differences in cell subset abundances between uninjured and burn-injured C57BL/6 SPF and NI mice. Neutrophils are circled in red, and B cells are circled in purple. (C, D) Statistical analysis of differences in blood immune cells in SPF and NI mice with and without burn injury. These CyTOF results are plotted as mean +/- SEM and are representative of 3 independent studies using 5 mice per group. Statistical analysis was assessed by two-way ANOVA with multiple comparisons *= p<0.05, **= p<0.005, ***= p<0.0005, ****= p<0.0001.

Plasma cytokine analysis revealed notable differences between SPF and NI mice at baseline and in response to burn injury (**Figure 3**). At baseline, NI mice exhibited elevated levels of IFNγ as compared to SPF mice.

**Figure 3.**
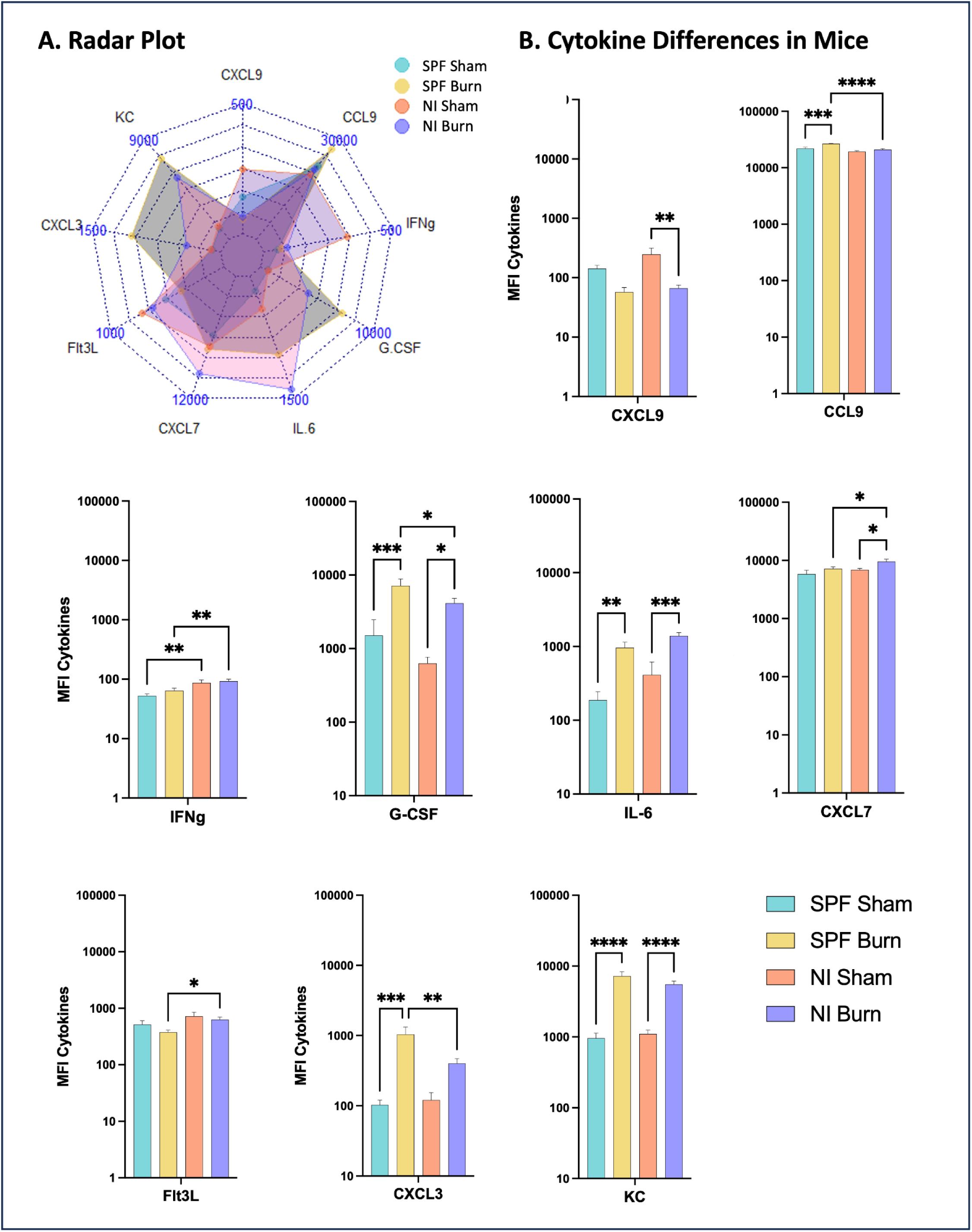
Plasma cytokine profiling using Luminex in Specific Pathogen-Free (SPF) and Natural Immune (NI) mice with and without burn injury. Plasma cytokine levels were measured using a Luminex assay. (A) Radar plot highlights differences in significant cytokines between SPF and NI mice with and without burn injury. (B) Bar graphs show individual cytokine MFI across SPF, SPF burn, NI, and NI burn groups. These Luminex results are plotted as mean +/- SEM and are representative of 3 independent studies using 5 mice per group. Statistical analysis was assessed by two-way ANOVA with multiple comparisons *= p<0.05, **= p<0.005, ***= p<0.0005, ****= p<0.0001. MFI = Mean Fluorescence Intensity.

Following burn injury, both SPF and NI mice demonstrated increased levels of G-CSF, IL-6, and KC. However, further cytokine responses diverged between groups—NI mice uniquely showed a burn-induced increase in CXCL7 and a decrease in CXCL9, while SPF mice exhibited increased levels of CCL9 and CXCL3. NI mice with burn injury had overall higher levels of IFNγ, CXCL7, and Flt3L, and lower levels of G-CSF and CXCL3 relative to SPF mice.

Analysis of bone marrow hematopoietic cells by flow cytometry demonstrated that both SPF and NI mice showed significant injury-induced increases in LK, MEP, and CLP (**Figure 4**). SPF mice had significant decreases in LT-HSC, while NI mice showed significant increases in the later progenitor populations CMP and GMP that were not seen in SPF mice (**Figure 4)**.

**Figure 4.**
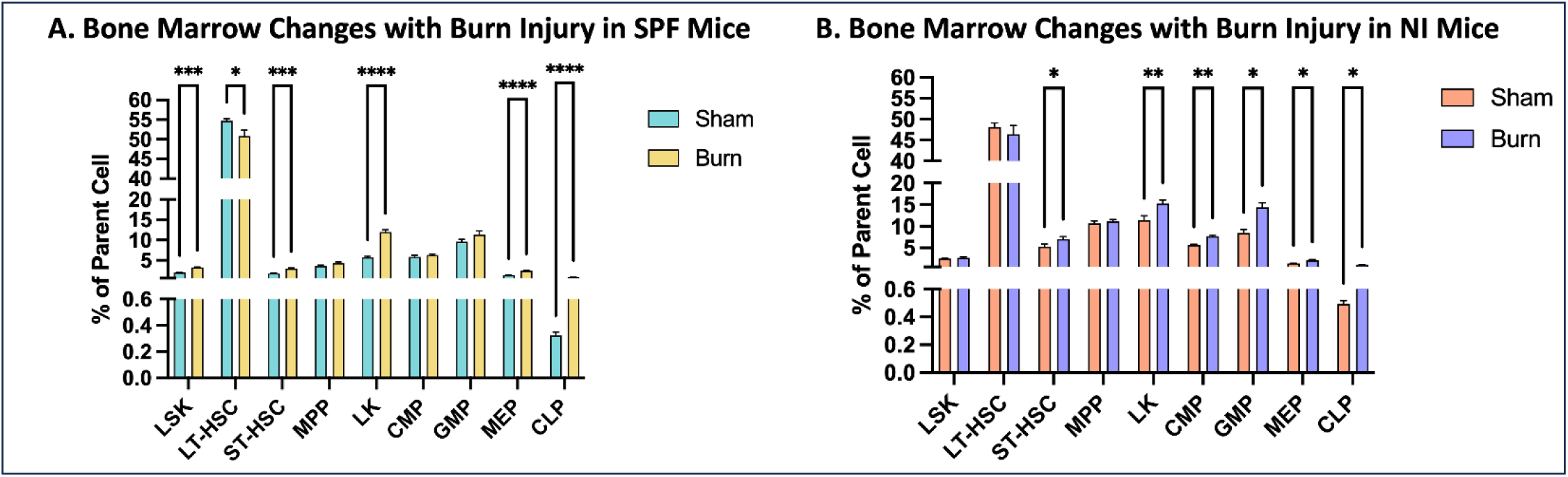
Flow Cytometry analysis comparing effects of burn injury on abundances of bone marrow cell populations in Specific Pathogen-Free (SPF) and Natural Immune (NI) mice. Bone marrow cells were stained by flow cytometry to detect hematopoietic stem and progenitor cells in uninjured and burn-injured C57BL/6 (A) SPF and (B) NI mice. These flow cytometry staining results are plotted as mean +/- SEM and are representative of 3 independent studies using 5 mice per group. Statistical analysis was assessed by two-way ANOVA with multiple comparisons *= p<0.05, **= p<0.005, ***= p<0.0005, ****= p<0.0001. LSK=Lineage(-)Sca-1(+)c- Kit(-) Cells; LT-HSC=Long-Term Repopulating Hematopoietic Stem Cells; ST-HSC=Short-Term Repopulating Hematopoietic Stem Cells; MPP=Multi-Potent Progenitor Cells; LK=Lineage(-)c-Kit(+)Sca-1(-); CMP=Common Myeloid Progenitor Cells; GMP=Granulocyte-Monocyte Progenitor Cells; MEP=Megakaryocyte-Erythrocyte Progenitor Cells; CLP=Common Lymphoid Progenitor Cells.

### Evaluating the two-hit trauma infection response in SPF and NI mouse models

To simulate a two-hit post-traumatic injury infection, mice underwent sham or burn injury, which was followed by an intranasal sublethal *P. aeruginosa* infection at 2 days after injury (**Figure 5A**). In the BM, two-hit infection stimulated decreases in LSK, LT-HSC, and CLP in SPF mice, with no significant changes in NI mice (**Figure 5B**). However, SPF mice demonstrated a significant increase in CD8^+^ T cells and a decrease in monocytes that did not occur in NI mice (**Figure 5C**). Both SPF and NI mice showed a decrease in B cells with burn injury and infection (**Figure 5C,D**). NI mice showed a large and significant increase in neutrophil percentage following the two-hit procedure that did not occur in SPF mice (**Figure 5D**).

**Figure 5.**
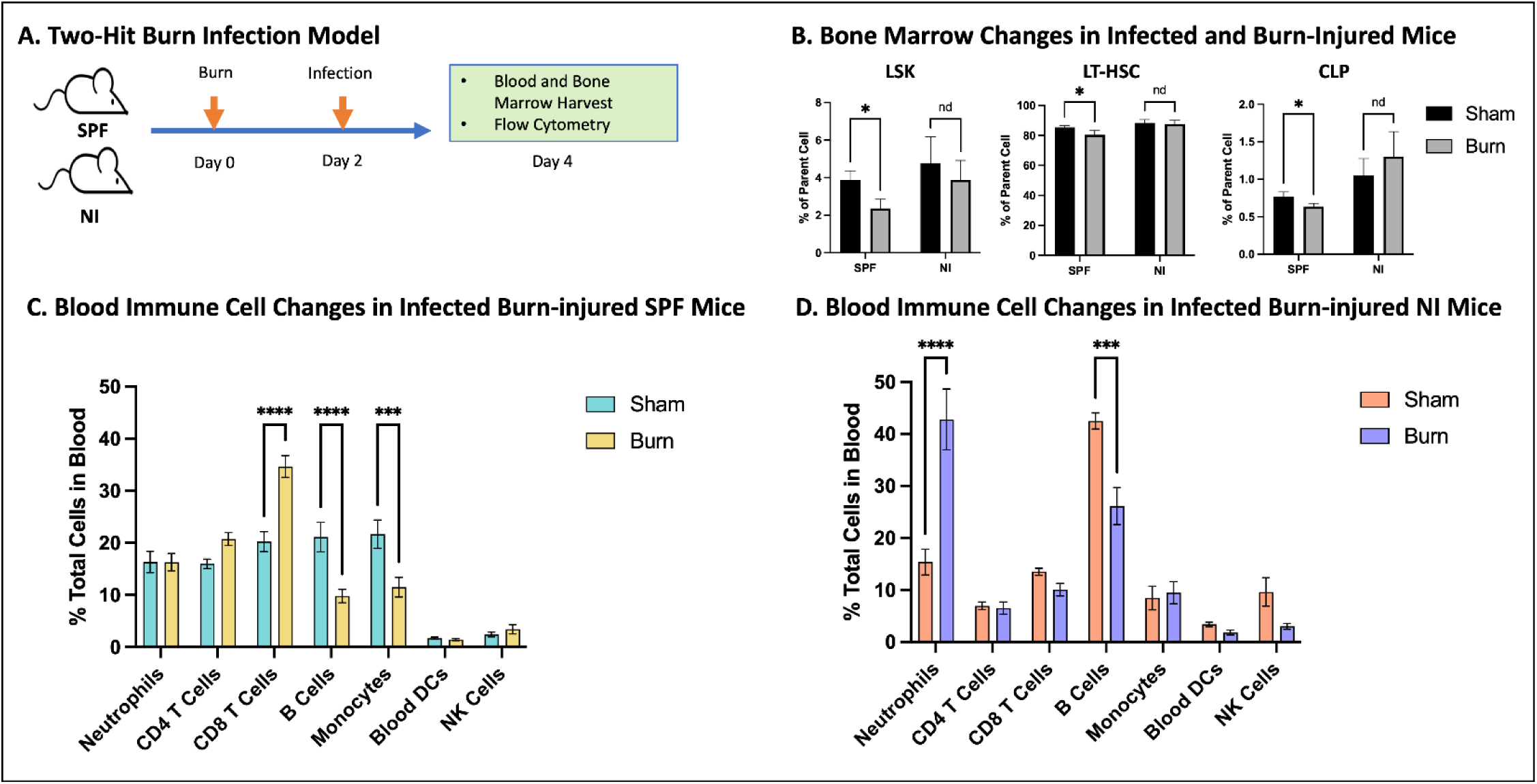
Comparing P. aeruginosa infection responses in Specific Pathogen-Free (SPF) and Natural Immune (NI) mice. (A) Plots comparing infection survival responses between SPF and NI mice (n=15 per group). (B) Total leukocytes at 48 hours post-infection were stained with a 48-marker CyTOF antibody panel to detect relative levels of the indicated immune cell subsets in infected and uninjured SPF and NI mice. (C,D) Bone marrow cells were stained by flow cytometry to detect hematopoietic stem and progenitor cells in infected and uninjured SPF and NI mice also at 48 hours. These CyTOF and flow cytometry staining results are plotted as mean +/- SEM and are representative of 3 independent studies using 5 mice per group. Statistical analysis was assessed by two-way ANOVA with multiple comparisons *= p<0.05, **= p<0.005, ***= p<0.0005, ****= p<0.0001. LSK=Lineage(-)Sca-1(+)c-Kit(-) Cells; LT-HSC=Long-Term Repopulating Hematopoietic Stem Cells; ST-HSC=Short-Term Repopulating Hematopoietic Stem Cells; MPP=Multi-Potent Progenitor Cells; LK=Lineage(-)c-Kit(+)Sca-1(-); CMP=Common Myeloid Progenitor Cells; GMP=Granulocyte-Monocyte Progenitor Cells; MEP=Megakaryocyte-Erythrocyte Progenitor Cells; CLP=Common Lymphoid Progenitor Cells.

There was no significant difference in *P. aeruginosa* infection survival between SPF and NI mice (**Figure 6A**). To assess the impact of infection alone on peripheral blood immune and bone marrow cells, mice were harvested 48 hours after infection for blood and bone marrow analysis by CyTOF and flow cytometry (**Figure 6B**). In the blood, SPF mice showed significant infection-induced increases in neutrophils and monocytes and decreases in B cells and NK cells (**Figure 6C**). In contrast, NI mice maintained stable neutrophil levels but exhibited significantly increased B cells and reduced CD4⁺ T cells and monocytes in response to infection (**Figure 6D**). In the bone marrow, infected SPF and NI mice both showed significant increases in LSK, LT-HSC, LK, and CLP cells and decreases in GMP and MEP cells (**Figure 6E,F**). Interestingly, SPF and NI mice had converse changes in ST-HSC and MPP, with an increase in SPF mice with infection, and a decrease in NI mice with infection (**Figure 6E,F**).

**Figure 6.**
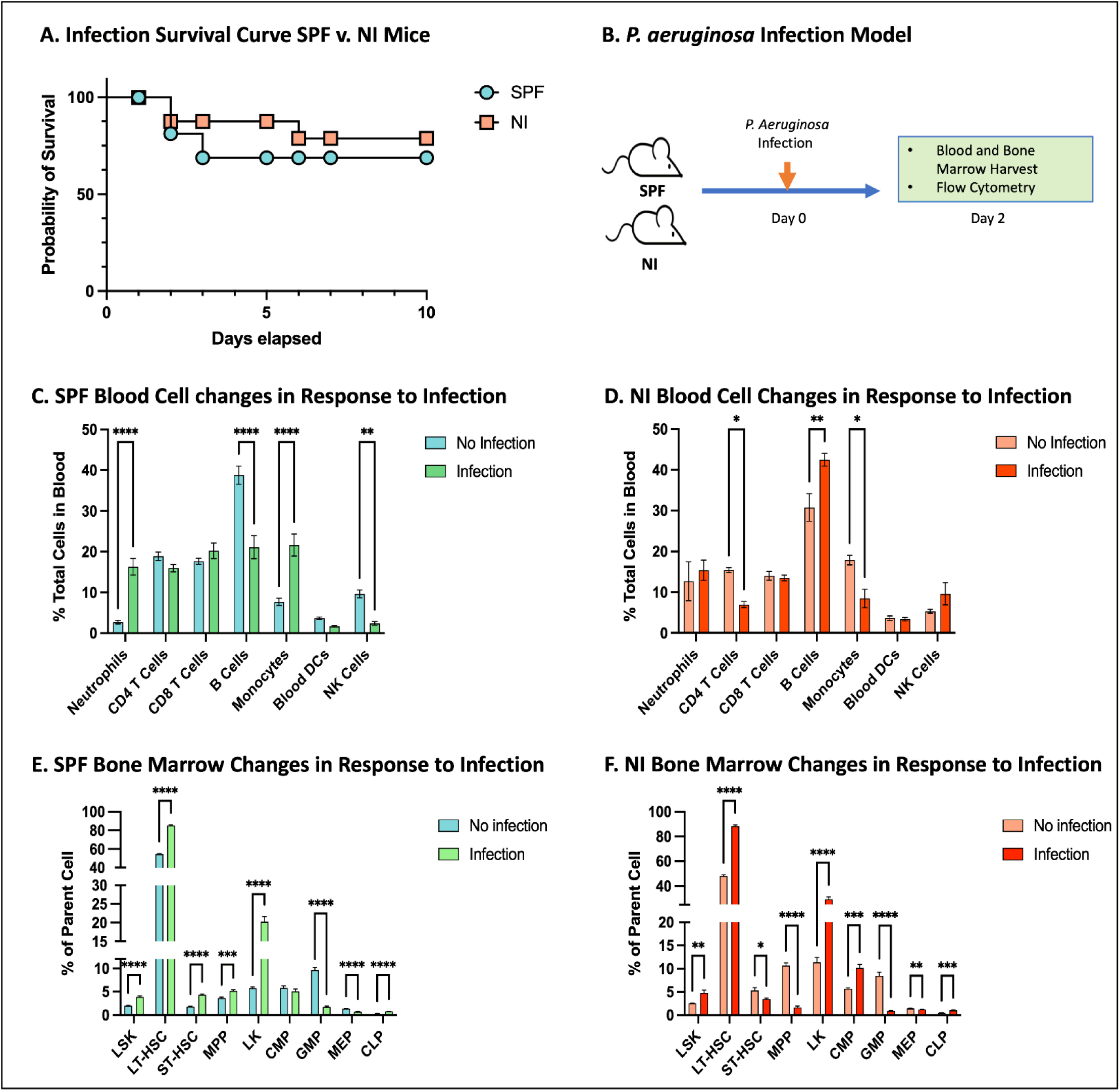
CyTOF and flow cytometry analysis comparing 2-hit infection blood and bone marrow cell responses in Specific Pathogen-Free (SPF) and Natural Immune (NI) mice. (A) Two-hit burn infection model workflow. (C,D) Total leukocytes were stained with a 48-marker CyTOF antibody panel to detect relative levels of the indicated immune cell subsets in *S. pneumoniae* infected uninjured (sham) and burn-injured C57BL/6 SPF and NI mice. (B) Plots comparing significant changes in bone marrow cell populations from uninjured and burn-injured C57BL/6 SPF and NI mice. These CyTOF and flow cytometry staining results are plotted as mean +/- SEM and are representative of 3 independent studies using 5 mice per group. Statistical analysis was assessed by two-way ANOVA with multiple comparisons, *= p<0.05, **= p<0.005, ***= p<0.0005, ****= p<0.00005. LSK=Lineage(-)Sca-1(+)c-Kit(-) Cells; LT-HSC=Long-Term Repopulating Hematopoietic Stem Cells; CLP=Common Lymphoid Progenitor Cells.

### Alterations in human and mouse neutrophil and monocyte composition following traumatic injury

Blood samples from patients who suffered traumatic injury were collected at 3 days following injury. Patient ages ranged from 21-78 (x̄=42) and 70% were male. Injury mechanisms included burn injury >40% TBSA (n=4), motor vehicle collision (n=3), fall (n=2), and penetrating injury (n=1). Age- and sex-matched healthy control blood samples were collected and prepared using the same method for comparison. CyTOF analysis was performed using a 48-marker CyTOF antibody panel that detects similar blood immune cell populations to those in mouse studies (**Supplementary Table 1**). Trauma patients exhibited a significant increase in circulating neutrophils compared to healthy controls (**Figure 7A,C**). Neutrophil clustering patterns, visualized using UMAP density contour plots, revealed distinct distributions between the two groups (**Figure 7A**). To compare blood immune cell responses between humans and mice, we used a whole blood method that better preserves mouse neutrophils. NI mice showed a significant increase in neutrophils with injury as well as a shift in neutrophil phenotypes that resembled what was observed in human trauma patients, while the neutrophil response in SPF did not show a phenotype shift (**Figure 7B,C**).

**Figure 7.**
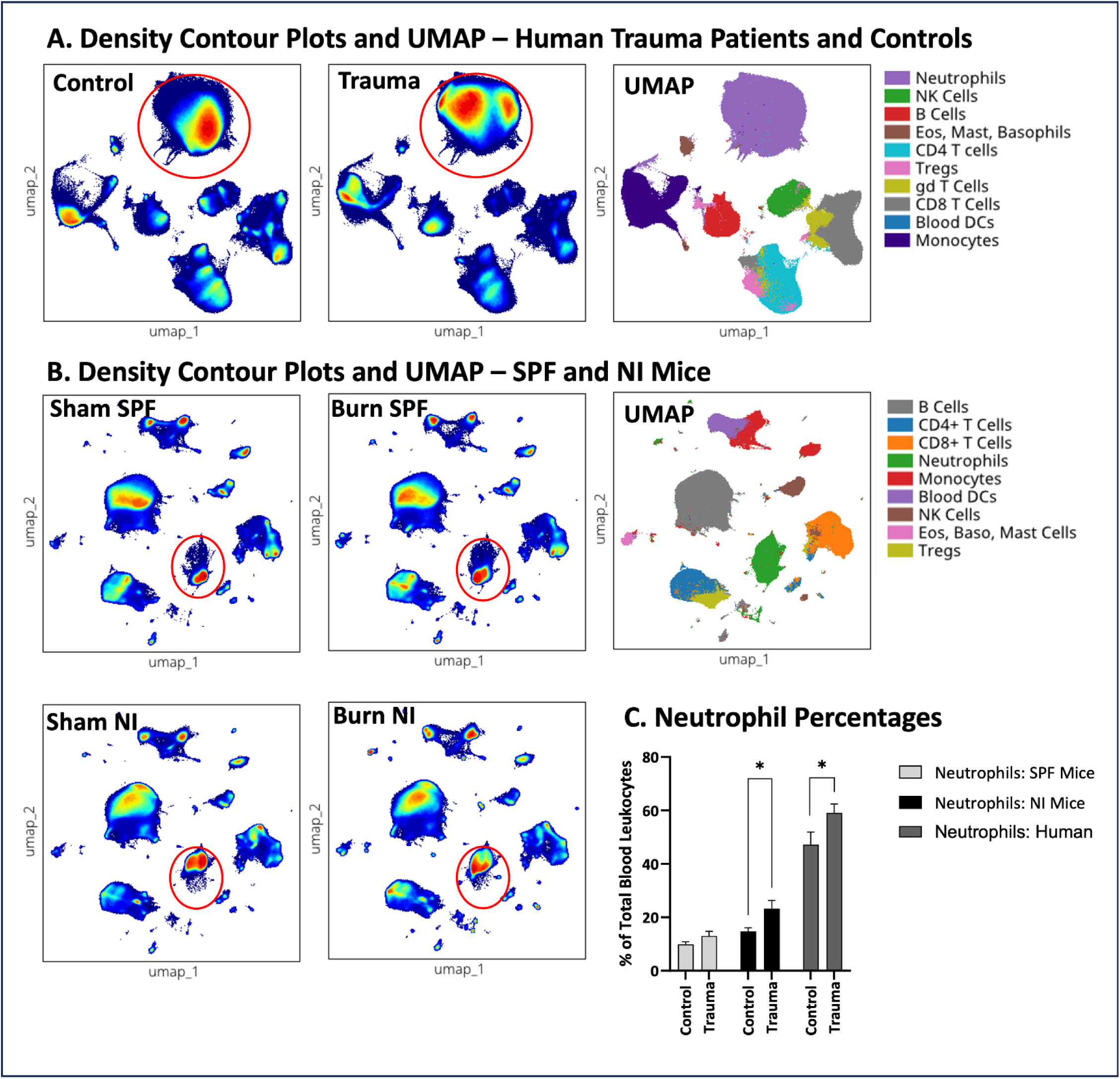
CyTOF analysis comparing effects of traumatic injury in humans, Specific Pathogen-Free (SPF), and Natural Immune (NI) mice on abundances of blood immune cells. (A) Total leukocytes were stained with a 48- marker CyTOF antibody panel to detect relative levels of the indicated immune cell subsets in healthy control patients (n=10) and day 3 trauma patients (n=10). Immune cells were clustered by markers into UMAP contour density heat plots for visualization. (B) SPF and NI whole blood was stained with a 48-marker CyTOF antibody panel to detect relative levels of the indicated immune cell subsets. These CyTOF staining results are plotted as density contour plots for visualization and neutrophils are circled in red (A,B). (C) Comparison of trauma-induced neutrophil abundance changes in SPF mice, NI mice, and humans showing controls vs. trauma for each species. These CyTOF results are plotted as mean +/- SEM and are representative of 3 independent studies using 5 mice per group. Statistical analysis was assessed by two-way ANOVA with multiple comparisons, *= p<0.05.

Comparison of neutrophil phenotypes between trauma patients and healthy controls revealed differential abundance of specific cell clusters (**Figure 8A,B**). Trauma patients exhibited a significantly increased proportion of flowSOM cluster (fc) fc_07 relative to controls, while the abundance of fc_10 remained comparable between the two groups (**Figure 8B**). Phenotypic characterization showed that fc_07, termed fresh or immature neutrophils demonstrated lower expression of CD15, CD16, and CD11b compared to fc_10, termed fully differentiated or mature neutrophils (**Figure 8B**). Trauma patients also demonstrated distinct alterations in monocyte expression patterns, further differentiating their immune profile from that of healthy controls. Four unique monocyte clusters were identified (fc_01, fc_02, fc_03, and fc_06), two of which showed significant differences between trauma and control patients (fc_01 and fc_06) (**Figure 8B**). Trauma patients had a significantly higher abundance of fc_06 and a lower abundance of fc_01 compared to controls. Cluster fc_06, termed fresh or immature monocytes, was characterized by reduced expression of CD172ab, CCR2, HO-1, HLA-DR, and CD11b relative to fc_01, termed mature monocytes (**Figure 8B**). Additional cluster analysis is shown in **Supplementary Figure 3**.

**Figure 8.**
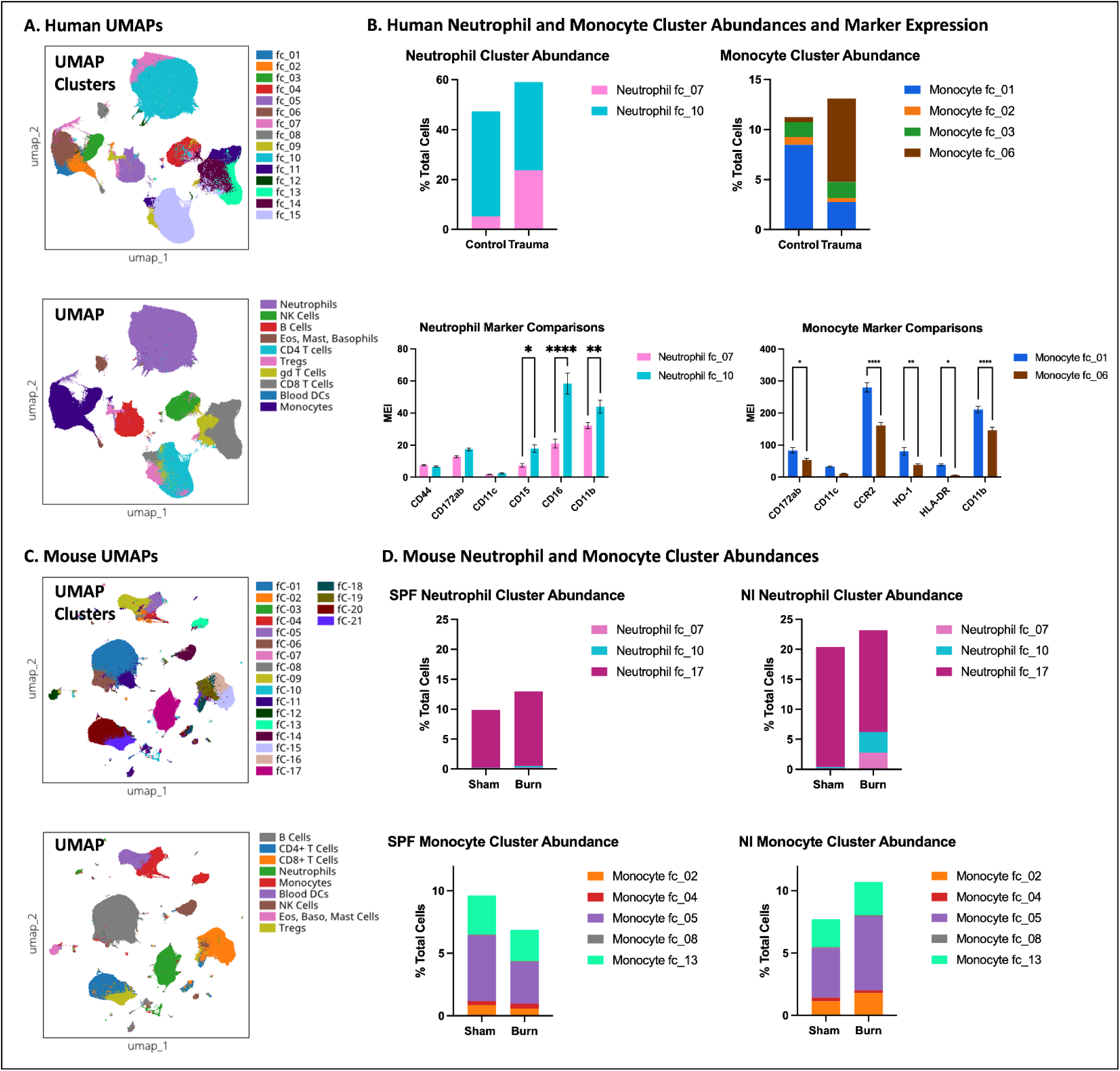
CyTOF analysis comparing effects of traumatic injury in humans, Specific Pathogen-Free (SPF), and Natural Immune (NI) mice on abundances of blood immune cells clusters and marker expression. (A) Total leukocytes were stained with a 48-marker CyTOF antibody panel to detect relative levels of the indicated immune cell subsets in healthy control patients (n=10) and day 3 trauma patients (n=10). Immune cells were clustered by markers into UMAP FlowSOM cluster organization map and UMAP cell type annotation plot for visualization. (B) Abundances of neutrophil and monocyte clusters are demonstrated in bar graphs and significant markers for notable neutrophil and monocyte clusters are also demonstrated in bar graphs. These CyTOF results are plotted as mean +/- SEM and are representative of 3 independent studies using 5 mice per group. Statistical analysis was assessed by two-way ANOVA with multiple comparisons *= p<0.05, **= p<0.005, ***= p<0.0005, ****= p<0.0001. MEI = Median Expression Intensity. (C) SPF and NI whole blood was stained with a 48-marker CyTOF antibody panel to detect relative levels of the indicated immune cell subsets. Immune cells were clustered by markers into UMAP FlowSOM cluster organization map and UMAP cell type annotation plot for visualization. Abundances of neutrophil and monocyte clusters are demonstrated in bar graphs for SPF and NI mice.

In mice, three distinct neutrophil clusters were identified (fc_07, fc_10, and fc_17) (**Figure 8C,D**). Following burn injury, NI mice demonstrated a significant increase in the abundance of clusters fc_10 and fc_17, a pattern not observed in SPF mice (**Figure 8D**). Cluster fc_10, termed fresh or immature proliferative neutrophils, was higher in TER 119, CD16/32, CD68, CD86 and lower in CD127 compared to clusters fc_07 and fc_17, and higher in Ki-67 compared to cluster fc_17 (**Supplementary Figure 4A,C-E**). Cluster fc_17, termed mature neutrophils, was higher in Ly6G and lower in CD16/32 and PU.1 compared to clusters fc_07 and fc_10. Monocyte analysis revealed five distinct clusters (fc_02, fc_04, fc_05, fc_08, and fc_13) present in both SPF and NI mice (**Figure 8D**). The abundance of cluster fc_05 decreased with burn injury in SPF mice but increased significantly in NI mice. Cluster fc_02 increased with burn injury in NI mice. In both SPF and NI mice, cluster fc_05, termed fresh or immature proliferative monocytes, was notable for high expression of Ly6C and Ki-67, with moderate levels of CD68, CD86, CCR2, and CD11b, and relatively low expression of CCR1, Sca-1, and I-A/I-E compared to other clusters (**Supplementary Figure 4B,F-J**). In contrast, cluster fc_02, termed mature, moderately activated monocytes, exhibited moderately elevated levels of CCR1, Ly6C, CD68, Sca-1, Ki-67, CD86, and I-A/I-E, but had lower CD11b expression relative to most other clusters.

## Discussion

Trauma remains a leading cause of morbidity and mortality for individuals in the United States, with a well-recognized immune dysregulation that predisposes these patients to subsequent infections. This increased susceptibility is thought to be attributed to the “two-hit response,” in which there is an initial inflammatory response to the traumatic insult followed by a counter-inflammatory response that leads to systemic immune suppression and increased susceptibility to secondary infections.^10,25,26^ Despite advancements in our understanding of post-trauma immune alterations, the underlying mechanisms remain complex and incompletely defined. One possible impediment is the limited physiological relevance of commonly used animal models, such as specific pathogen-free (SPF) mice, which do not fully capture the diversity of human immune cell composition or responses.

While numerous mouse models for traumatic injury have been developed to study these trauma-related immune changes, these models do not fully capture the immune complexity of human trauma patients for several reasons. First, inbred mice are often used to model human diseases to provide a controlled experimental platform by removing genetic variables. This approach is appropriate to model human disease mechanisms and basic mammalian biology but fails to capture the genetically diverse nature of human physiology. Second, laboratory mice were established to be SPF in the 1960s to further reduce microbial variability.^27,28^ However, this approach also affected normal development processes of the immune system. A definitive report in 2016 documented that SPF mice do not have fully developed immune system but can be converted to a more “human-like” immune system by co-housing with pet shop “dirty” mice that contribute normal flora to the environment.^15^ This observation has been confirmed by other investigators using re-wilding and co-housing approaches.^15,29,30^

We established a novel “dirty” mouse model by co-housing inbred female C57BL/6, Balb/c, or CD-1 mice with dirty mice followed by breeding these converted mice for multiple generations. We believe multigenerational mice have an advantage over the co-housed dirty mouse approaches by providing stable colonies of laboratory mice with natural and stable changes in their immune system. These multigenerational mice, which we call natural immune (NI) mice may better approximate human immune physiology due to their lifelong exposure to environmental commensals and pathogens. To further support this, we profiled microbial taxa in SPF and NI mice and found that NI mice harbor an entirely different community of commensal and environmental microbes compared to SPF mice, consistent with a history of sustained pathogen exposure (**Supplementary Figure 1**). Thus, these mice would logically provide a more translatable model for immunology research.

While the overall composition of the immune system in mice and humans is similar, there are significant differences in immune cell subset abundances. In humans, blood immune cells are predominantly 50-70% neutrophils, with lymphocytes comprising 20-50% of all peripheral blood immune cells. In contrast, mice exhibit a lymphocyte-dominant blood immune cell profile of 70-90% lymphocytes with only 10-20% neutrophils.^31,32^ A major observation from our study is that NI mice exhibit a baseline blood immune cell profile that more closely resembles humans, with higher percentages of circulating neutrophils and lower B cells than SPF mice (**Figure 1B**). This suggests that NI mice have a more developed myeloid-rich immune system, possibly due to natural microbial or pathogen exposure during their lifespan.^15^ This finding suggests that natural microbial exposure in NI mice promotes maturation of a more myeloid-rich immune system, possibly through sustained environmental immune stimulation. Such persistent exposure may induce long-term immune adaptation consistent with the concept of trained immunity. This is supported by the observation that NI mice have higher baseline progenitor cell populations in the bone marrow, suggesting a primed hematopoietic system that may be epigenetically or metabolically conditioned to maintain a heightened state of immune readiness.

These findings underscore the value of multigenerational “dirty” mouse models, as they appear to stably acquire traits indicative of trained immunity. In contrast, co-housed SPF mice would lack these durable adaptations, limiting their translational utility for modeling human immune responses after trauma or infection.

To investigate how this baseline conditioning influences the immune response to injury, we applied a well-established burn trauma model and compared peripheral and bone marrow immune changes in SPF and NI mice. We observed that SPF mice displayed a significant increase in neutrophils and monocytes, along with a decline in B cells and CD4+ T cells following injury (**Figure 2C,D**). In contrast, NI mice exhibited minimal changes in these populations **(Figure 2C,D)**. Given that NI mice have a higher baseline neutrophil and lower baseline B cell counts, this muted response likely reflects a preconditioned immune system rather than a blunted reaction to injury.

To assess systemic immune responses to injury, we measured plasma cytokines using a multiplexed Luminex assay approach. At baseline, NI mice had elevated IFNγ (**Figure 3**), a cytokine associated with immune activation and macrophage priming, suggesting a more proinflammatory immune system set point. Following burn injury, both groups showed increased G-CSF, IL-6, and KC—key mediators of neutrophil mobilization and the acute phase response **(Figure 3)**. However, distinct group-specific patterns emerged. NI mice upregulated CXCL7, a chemokine involved in platelet activation and tissue repair, and downregulated CXCL9, a T cell chemoattractant linked to chronic inflammation (**Figure 3**). In contrast, SPF mice exhibited increased levels of CCL9 and CXCL3, both associated with monocyte and neutrophil recruitment (**Figure 3**). Compared to SPF mice, burn-injured NI mice had higher IFNγ, CXCL7, and Flt3L (a cytokine supporting dendritic cell development), and lower levels of G-CSF and CXCL3 (**Figure 3**). These patterns suggest that NI mice mount a more targeted and regulated cytokine response to injury, potentially balancing inflammation with tissue repair. The altered cytokine profile in NI mice reflects a “trained” immune state that may confer greater readiness and adaptability to trauma, aligning more closely with immune responses observed in human injury.

Interestingly, burn-injured SPF mice demonstrated changes in early hematopoietic stem and progenitor cells, including significant decreases in LT-HSCs, while NI mice showed significant injury-induced increases in downstream progenitor populations, including CMPs and GMPs—changes that were not observed in SPF mice (**Figure 4**). Both SPF and NI mice exhibited increases in LK, MEP, and CLP populations following injury (**Figure 4**). This pattern suggests that NI mice mount a more rapid and directed hematopoietic response, potentially reflecting a more “primed” or “trained” immune state that favors efficient replenishment of innate immune cells in response to trauma.

Given the increased susceptibility of trauma patients to infections, we expanded our model by introducing a sublethal *Pseudomonas aeruginosa* lung infection following burn injury. This model mimics the two-hit response that has been described in trauma patients.^3,4,21^ First, we examined the impact of infection alone on blood and bone marrow cell changes. We found that despite no differences in survival between SPF and NI mice, their immune responses to infection were significantly different. Following infection, SPF mice showed increase neutrophil levels that were similar to those found in uninfected NI mice (**Figure 6C,D**). In contrast, NI mice did not show a significant increase in neutrophils following infection (**Figure 6D**). This may be because there is already sufficient blood neutrophil levels to react to lung *P. aeruginosa* infection. However, NI mice did show significantly more B cells as well as fewer CD4^+^ T cells and monocytes than SPF mice following infection (**Figure 6C,D**). This indicates major differences in infection responses between SPF and NI mice.

Results in the bone marrow revealed that both SPF and NI mice exhibited expansion of early progenitor populations (LSK, LT-HSC, LK, and CLP) and reductions in GMP and MEP populations following infection (**Figure 6E,F**). Notably, SPF and NI mice showed opposing responses in ST-HSC and MPP populations, with increases in SPF mice and decreases in NI mice, suggesting that NI mice may rely less on expansion of intermediate progenitors and instead exhibit a distinct hematopoietic adaptation to infection.

Second, following burn injury, SPF and NI mice exhibited wildly different immune responses to *P. aeruginosa* infection. Burn-injured NI mice mounted a dramatic neutrophil response (**Figure 5D**), suggesting a heightened capacity to combat secondary infections in this two-hit trauma with infection model. In contrast, burn-injured SPF mice did not show increases in blood neutrophils but did demonstrate increased CD8^+^ T cells and reduced monocytes (**Figure 5C**). These contrasting responses to two-hit infection support major differences in how injured SPF and NI mice respond to infections. We hypothesize that NI mice may better mimic the human two-hit response to infections. Both SPF and NI mice experienced a decline in B cells, reinforcing the impact of trauma and infection on adaptive immunity (**Figure 5C,D**). In the bone marrow, SPF mice exhibited significant depletion of hematopoietic stem and progenitor cells (LSK, LT-HSC, and CLP), whereas NI mice maintained stable progenitor populations, suggesting a more resilient hematopoietic response to infection following burn trauma (**Figure 5B**). Taken together, these findings indicate that NI mice possess a preconditioned or trained immune state that enables a more robust emergency granulopoiesis response to infection while preserving bone marrow function, which may better model human trauma two-hit infection responses.

To better contextualize our findings, we analyzed peripheral blood immune composition in human trauma patients. As expected, trauma patients exhibited a significant increase in circulating neutrophils compared to healthy controls, mirroring the increase observed in SPF mice and the baseline immune state of NI mice (**Figure 7A,C**). However, unlike SPF mice, NI mice did not require an injury-induced neutrophil increase to reach levels comparable to human trauma patients. Additionally, CyTOF analysis revealed distinct neutrophil clustering patterns between trauma patients and healthy controls (**Figure 7A, 8A,B**), with trauma patients showing an enrichment of a fresh or immature subset of neutrophils expressing lower levels of CD15, CD16, and CD11b, suggesting an expansion of immature neutrophils following trauma, consistent with emergency granulopoiesis or mobilization of immature neutrophils. This pattern was also observed in NI mice following injury but was not seen in SPF mice (**Figure 7B, 8D**). In mice, CyTOF analysis revealed three neutrophil clusters (fc_07, fc_10, and fc_17) (**Figure 8D**). NI mice demonstrated a post-injury expansion of fresh or immature and proliferative neutrophils (**Figure 8D, Supplementary Figure 4**), not seen in SPF mice, mirroring the human trauma neutrophil response (**Supplementary Figure 4**). These neutrophil subsets were marked by high TER119, CD16/32, CD68, CD86, and Ki-67, and low CD127 (**Supplementary Figure 4**), suggestive of proliferative and recently emigrated bone marrow neutrophils. These findings parallel the expansion of low CD15, CD16, and CD11b neutrophils in human trauma, pointing to conserved mechanisms of emergency granulopoiesis between NI mice and humans.

While both species demonstrated injury-induced remodeling of monocyte populations, the direction and nature of these shifts differed. In humans, trauma was associated with an increase in fresh or immature monocytes, a subset defined by low expression of CD172ab, CCR2, HO-1, HLA-DR, and CD11b (**Figure 8A, Supplementary Figure 3**). Conversely, mature monocytes were reduced in abundance in trauma patients. These findings suggest trauma skews monocyte composition towards an immature phenotype, which may contribute to the well-documented immune dysfunction seen in critically injured patients. In contrast, mouse monocyte responses were highly dependent on prior pathogen exposure. In SPF mice, the inflammatory monocyte subset fc_05 (high Ly6C, Ki-67) decreased after burn injury, while in NI mice it increased significantly (**Figure 8D, Supplementary Figure 4**). Similarly, fc_02, a moderately activated monocyte population, expanded in NI mice but not in SPF counterparts (**Figure 8D, Supplementary Figure 4**).

Compared to humans, mice showed a broader range of activation states. While both humans and NI mice demonstrated expansion of non-classical or regulatory-like monocyte subsets post-injury, SPF mice failed to mirror these changes, reinforcing the importance of using microbiota-exposed dirty mouse models for translational relevance.

A central aim of this study was to compare immune system phenotypes between SPF and NI mice. The observations that the preconditioned trained immune state in NI mice, their hematopoietic shift toward later progenitor expansion, and their more robust secondary infection response all suggest that lifelong exposure to natural environmental commensal microbes or pathogens greatly influences the baseline immune system and responses to injury and infection. NI mice may thus serve as a more physiologically relevant trauma immunology research animal model. In addition, these mice may serve as founder mice for others that are interested in using NI mice in their research programs. These findings underscore the need for further research into the epigenetic modifications, innate immune memory, and trained immunity mechanisms in these mouse models, which was outside the scope of the present study. By leveraging NI mice, we may be able to develop more predictive preclinical models that better inform therapeutic strategies for trauma patients, particularly in the context of immune modulation and infection management.

## Supporting information

Featured Image

## Abbreviations

CD: Cluster of Differentiation
CD4+ T cells: CD4+ T Lymphocytes
CD8+ T cells: CD8+ T Lymphocytes
CLP: Common Lymphoid Progenitors
CMP: Common Myeloid Progenitors
CyTOF: Cytometry by Time-of-Flight
GMP: Granulocyte-Macrophage Progenitors
HLA-DR: Human Leukocyte Antigen - DR isotype
HSPC: Hematopoietic Stem and Progenitor Cells
LK: Lineage-Kit (Lineage-negative, c-Kit-positive)
LSK: Lineage-SCM-Kit (Lineage-negative, Sca-1-positive, c-Kit-positive)
LT-HSC: Long-Term Hematopoietic Stem Cells
MPP: Multipotent Progenitors
NI: Natural Immune
NK cells: Natural Killer Cells
P. aeruginosa: *Pseudomonas aeruginosa*
RBC: Red Blood Cells
SPF: Specific Pathogen-Free
ST-HSC: Short-Term Hematopoietic Stem Cells
UMAP: Uniform Manifold Approximation and Projection

## Authorship

**AB** – Project conceptualization; Investigation – mouse burn injury and infection; Data curation – flow cytometry and CyTOF staining; Formal analysis – flow cytometry and CyTOF; Writing – original draft, review, editing, and revision.

**JP** – Investigation – mouse burn injury and infection; Data curation – flow cytometry and CyTOF staining; Formal analysis – flow cytometry and CyTOF; Writing – review, editing, and revision.

**EM** – Investigation – mouse burn injury and infection; Data curation – flow cytometry, CyTOF staining, and Luminex assays; Formal analysis – flow cytometry, CyTOF, and Luminex.

**ZM** – Investigation – mouse burn injury and infection; Writing – review, editing, and revision.

**CL** – Investigation – mouse burn injury and infection; Writing – review, editing, and revision.

**BN** – Investigation – mouse burn injury and infection; Data curation – flow cytometry and CyTOF staining.

**AS** – Data curation – trauma and control patient datasets. Writing – review, editing, and revision. **GB** – Data curation – trauma and control patient datasets. Writing – review, editing, and revision. **AZ** – Natural immune mouse colony maintenance.

**FB** – Conceptualization – natural immune mouse colony formation; Writing – review, editing, and revision. **JL** – Project conceptualization; Natural immune mouse colony formation; Investigation – mouse burn injury and infection; Data curation – flow cytometry and CyTOF staining; Formal analysis – flow cytometry and CyTOF; Writing – original draft, review, editing, and revision.

## Acknowledgments

We gratefully acknowledge the patients who generously provided informed consent for blood sample collection. Their contributions were essential to the success of this study and to advancing our understanding of the immune response to trauma. Funded by NIH R01AI177803, R01AI148232, P30AR070253, and the Gillian Reny Stepping Strong Center for Trauma Innovation (SSCTI). Graphic abstract was created with BioRender.^33^

