## Supplementary Data and Legends for "A Multigenerational “Dirty” Mouse Model for Studying Trauma-Induced Immune Dysregulation and Infection Susceptibility"

#### Supplementary Figures and Table

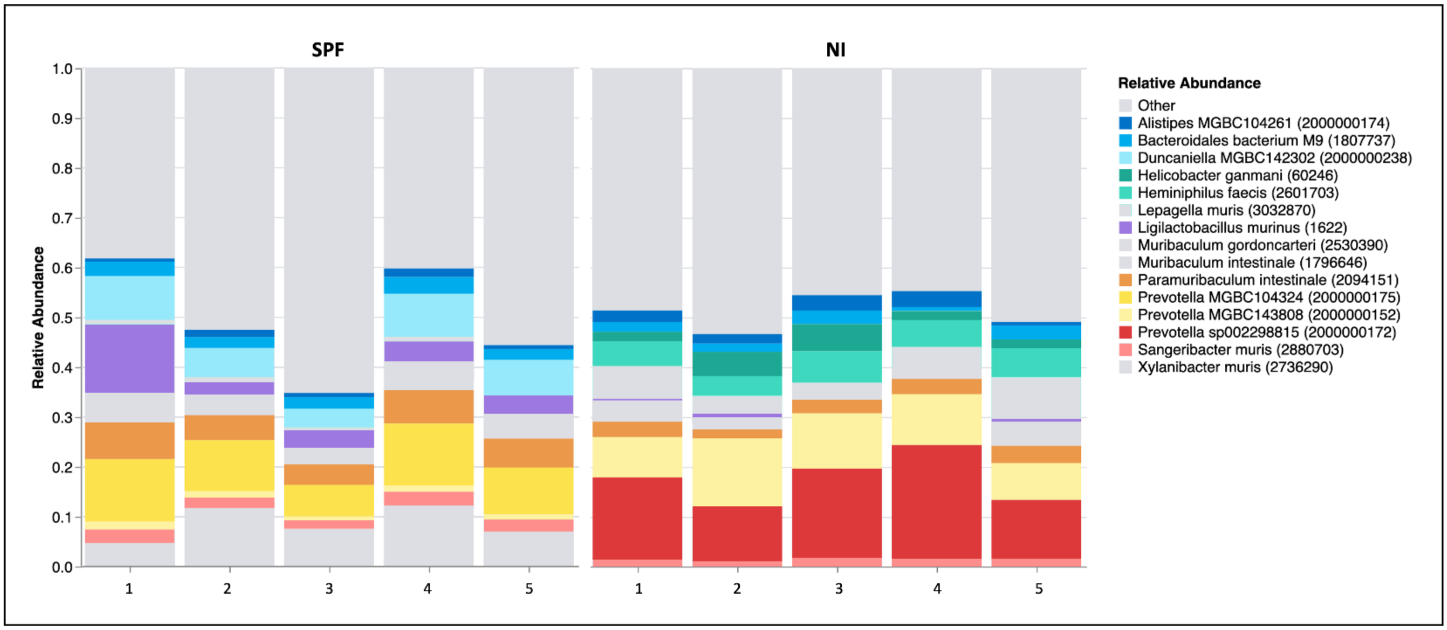


**Supplementary Figure 1.** *Gut Microbial Profiles in specific pathogen-free (SPF) and natural immune (NI) Mice.* Relative abundance of gut microbial taxa in stool samples from SPF and NI mice, analyzed using the Transnetyx metagenomics platform with One Codex profiling (Cordova, TN). Each bar represents an individual mouse (n = 5 per group), and the data are representative of three independent experiments.

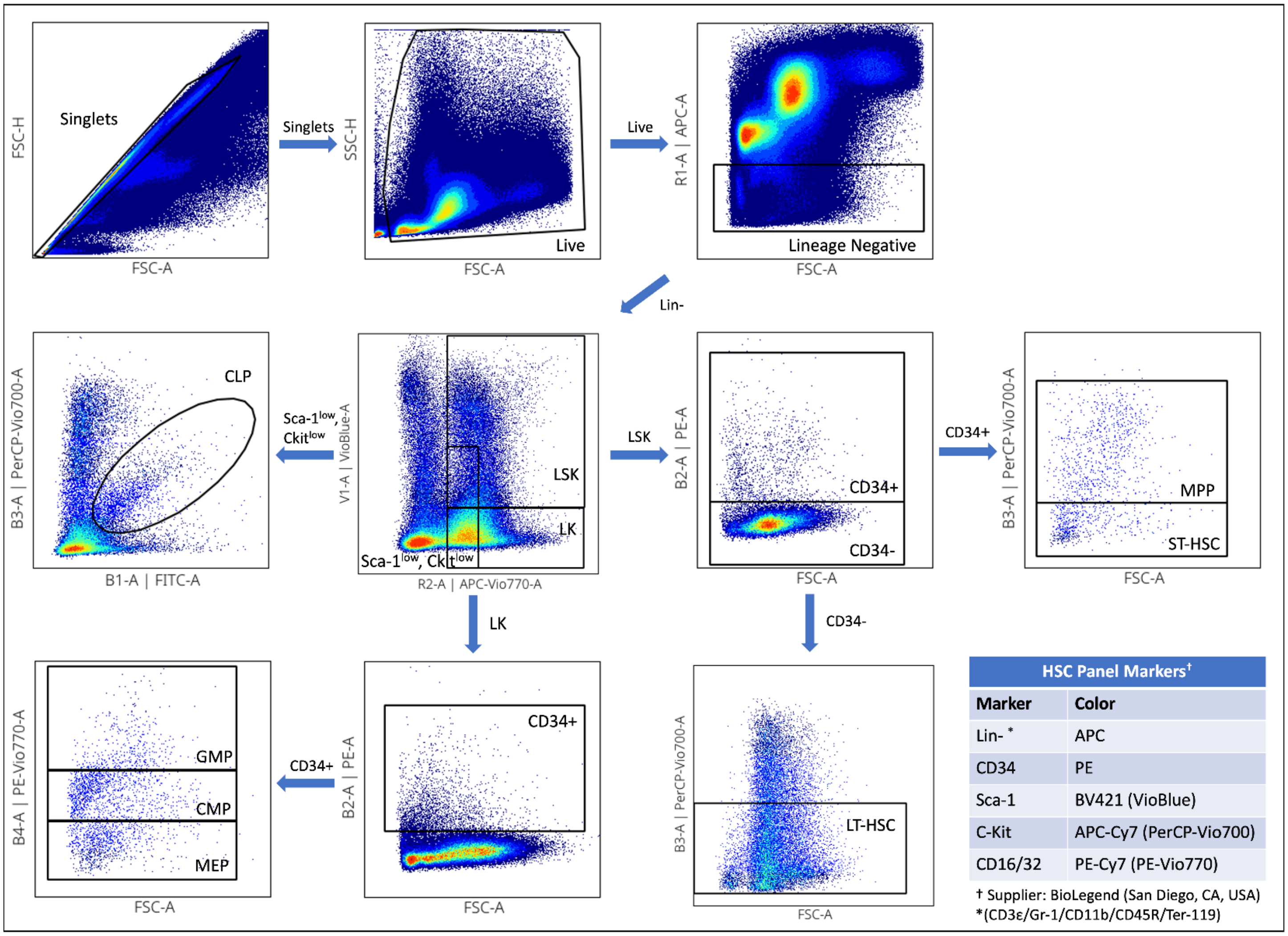


| HSC Panel Markers† | |
|---|---|
| **Marker** | **Color** |
| Lin- * | APC |
| CD34 | PE |
| Sca-1 | BV421 (VioBlue) |
| C-Kit | APC-Cy7 (PerCP-Vio700) |
| CD16/32 | PE-Cy7 (PE-Vio770) |

† Supplier: BioLegend (San Diego, CA, USA)
*(CD3ε/Gr-1/CD11b/CD45R/Ter-119)

**Supplementary Figure 2.** *Hematopoietic Stem Cell (HSC) gating scheme.* The table insert lists the fluorescent-tagged antibodies or cocktails used to gate specific bone marrow stem and progenitor cell populations. The diagram shows the gating steps for each indicated population. From these gates, we generated the percentages of cells within each stem or progenitor cell populations (see Figure 1).

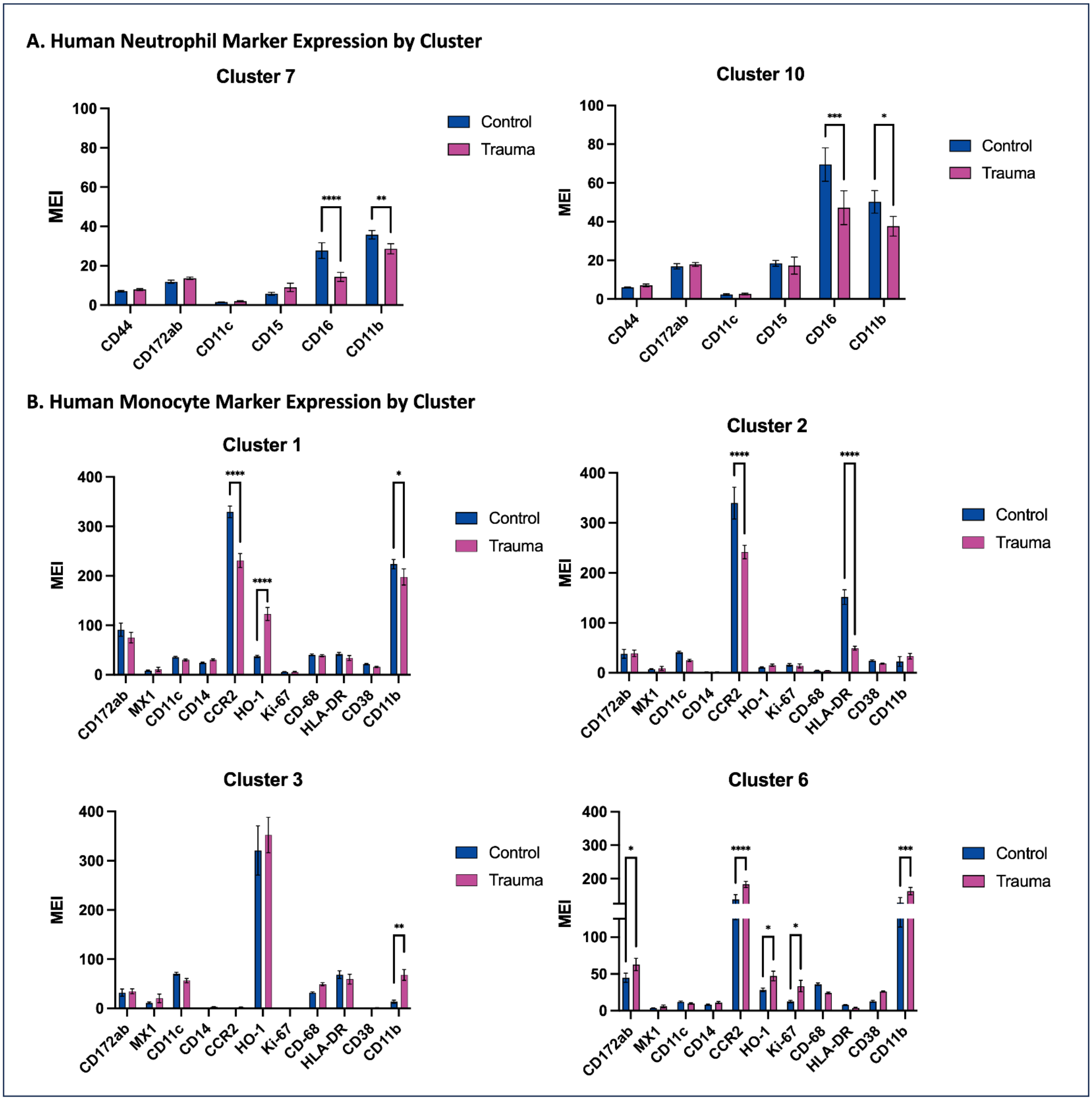


**Supplementary Figure 3.** *Human neutrophil and monocyte marker expression by clusters.* Total leukocytes were stained with a 48-marker CyTOF antibody panel to detect relative levels of the indicated immune cell subsets in healthy control patients (n=10) and day 3 trauma patients (n=10). Immune cells were clustered by markers into UMAP FlowSOM clusters. (A) Neutrophil clusters (fc_07, fc_10) were identified and markers of interest are presented in bar graphs. (B) Monocyte clusters (fc_01, fc_02, fc_03, fc_06) were identified and markers of interest are presented in bar graphs. Statistical analysis was assessed by two-way ANOVA with multiple comparisons *= p<0.05, **= p<0.005, ***= p<0.0005, ****= p<0.0001. MEI = Median Expression Intensity.

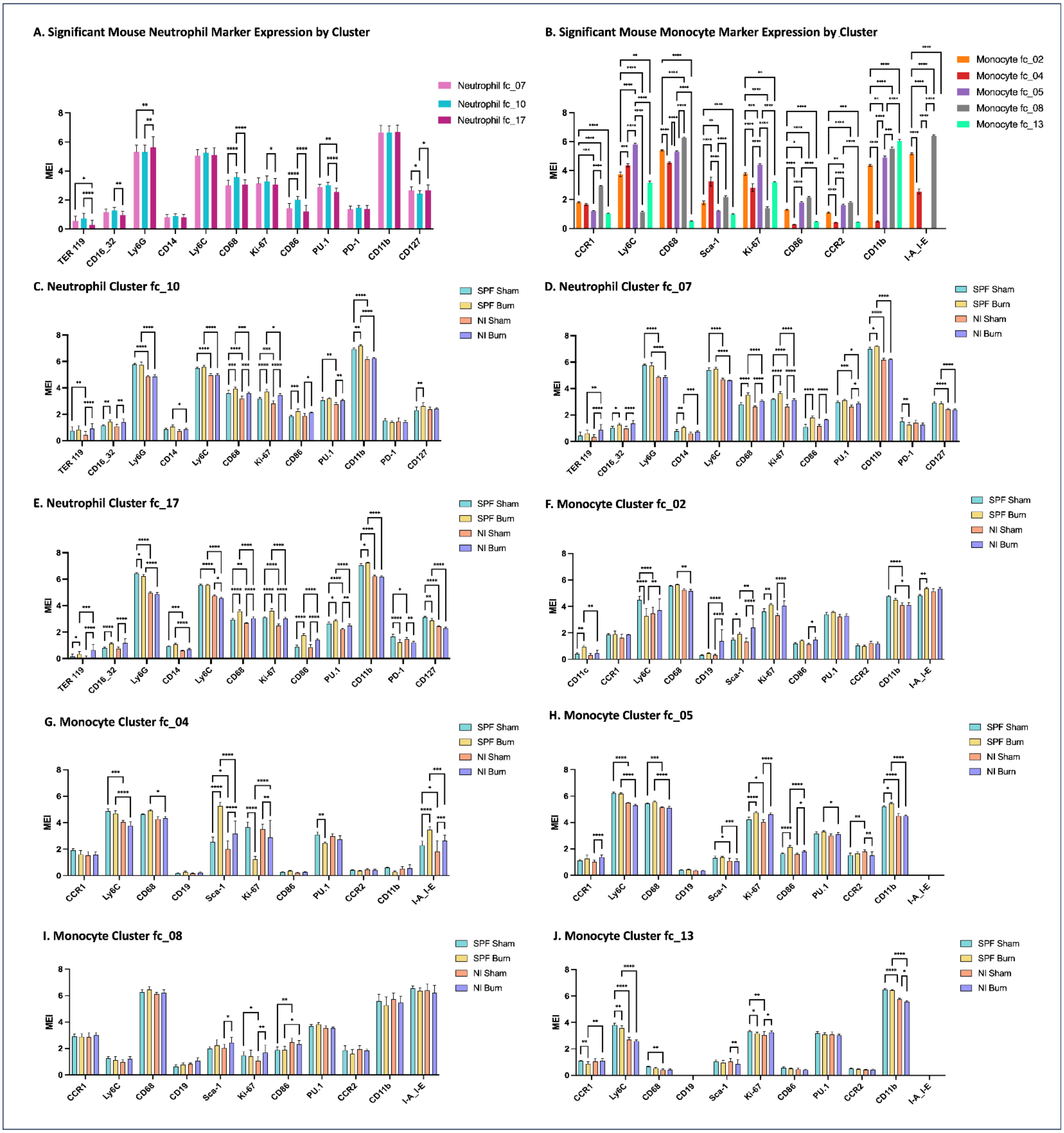


**Supplementary Figure 4.** *Mouse neutrophil and monocyte marker expression by cluster.* Specific Pathogen-Free (SPF) and Natural Immune (NI) mouse whole blood was stained with a 48-marker CyTOF antibody panel to assess immune cell phenotypes following burn injury. (A) Significant marker expression for neutrophil clusters (fc_07, fc_10, fc_17). (B) Significant marker expression for monocyte clusters (fc_02, fc_04, fc_05, fc_08, fc_13). (C–E) Expression of markers of interest for individual neutrophil clusters. (F–J) Expression of markers of interest for individual monocyte clusters. Statistical analysis was performed using two-way ANOVA with multiple comparisons. *p<0.05, **p<0.005, ***p<0.0005, ****p<0.0001. MEI = Median Expression Intensity.

**Supplementary Table 1.** *Mouse and Human CyTOF Phenotyping Antibody Panel.*

**A. Mouse CyTOF Phenotyping Antibody Panel**

| Marker | Clone | Metal | Supplier |
|---|---|---|---|
| Arginase | polyclonal | 89Y | * |
| B220/CD45R | RA3-6B2 | 111Cd | † |
| TER-119 | Erythroid Cells | 112Cd | † |
| CD16/32 | 93 | 113Cd | † |
| CD4 | RM4-5 | 114Cd | † |
| CD44 | IM7 | 115In | † |
| CD8a | 53-6.7 | 116Cd | † |
| CD45 | 30-F11 | 141Pr | † |
| ASC/TMS1/PYCARD | D2W8U | 142Nd | ‡ |
| CD49b | DX5 | 143Nd | † |
| CD184 (CXCR4) | L276F12 | 144Nd | † |
| CD314 (NKG2D) | CX5 | 145Nd | † |
| CD11c | N418 | 146Nd | † |
| CD223 (LAG3) | C9B7W | 147Sm | † |
| Ly6G | 1A8 | 148Nd | † |
| CCR1 | 643854 | 149Nd | * |
| CD14 | Sa14-2 | 150Nd | † |
| Ly6C | HK1.4 | 151Eu | † |
| CD3 | 154-2C11 | 152Sm | † |
| CD152 (CTLA4) | UC10-4B9 | 153Eu | † |
| CD103 | 2E7 | 154Sm | † |
| CD68 | FA-11 | 155Gd | † |
| CD19 | 6D5 | 156Gd | † |
| CD186 (CXCR6) | SA051D1 | 157Gd | † |
| T-bet | 4B10 | 158Gd | † |
| CD206 (MMR) | C068C2 | 159Tb | † |
| Sca-1 (Ly-6A-E) | E13-161.7 | 160Gd | † |
| CD83 | Michel-19 | 161Dy | † |
| FoxP3 | FJK-16s | 162Dy | § |
| NK1.1 | PK136 | 163Dy | † |
| Ki-67 | 8D5 | 164Dy | † |
| CD115 | 460615 | 165Ho | * |
| CD86 (B7-2) | GL-1 | 166Er | † |
| CD25 | 3C7 | 167Er | † |
| CD117(c-kit) | 2B8 | 168Er | † |
| PU.1 | phpu13 | 169Tm | † |
| CD278 (ICOS) | C398.4A | 170Er | † |
| CD279 (PD-1) | 29F.1A12 | 171Yb | † |
| CD192(CCR2) | QA18A56 | 172Yb | † |
| CD69 | H1.2F3 | 173Yb | † |
| CD11b | M1/70 | 174Yb | † |
| F4/80 | BM8 | 175Lu | † |
| IL-23R | 12B2B64 | 176Yb | † |
| CD127 | A7R34 | 194Pt | † |
| CD5 | 53-7.3 | 195Pt | † |
| Caspase-1 | 5B10 | 196Pt | † |
| CD34 | MEC14.7 | 198Pt | † |
| I-A/I-E (MHC-II) | M5/114.15.2 | 209Bi | † |

*R&D Systems (Minneapolis, MN, USA)
†BioLegend (San Diego, CA, USA)
‡Cell Signaling Technology (Danvers, MA, USA)
§eBioscience (San Diego, CA, USA)

**B. Human CyTOF Phenotyping Antibody Panel**

| Marker | Clone | Metal | Supplier |
|---|---|---|---|
| CD44 | IM7 | 089Y | † |
| CD172ab | SE5A5 | 111Cd | † |
| CD8a | RPA T8 | 112Cd | † |
| CD20 | 2H7 | 113Cd | ¶ |
| CD4 | RPA T4 | 114Cd | † |
| CD3 | UCHT1 | 115In | † |
| CD56 | NCAM16.2 | 116Cd | \|\| |
| CD45 | HI30 | 141Pr | † |
| TLR4 | 610015 | 142Nd | * |
| MX1 | D3W7I | 143Nd | ‡ |
| CD64 | 10.1 | 144Nd | † |
| PU.1 | phpu13 | 145Nd | § |
| C3AR | hC3aRZ8 | 146Nd | † |
| CD45RO | REA611 | 147Sm | † |
| GZMA | CB9 | 148Nd | † |
| GZMK | GM26E7 | 149Sm | † |
| CD11c | Bu15 | 150Nd | † |
| CD123 | 6H6 | 151Eu | † |
| CD14 | M5E2 | 152Sm | † |
| CD69 | FN50 | 153Eu | † |
| CD15 | MC-480 | 154Sm | † |
| Siglec-1 | 7-239 | 155Gd | † |
| CD8b | SIDI8BEE | 156Gd | § |
| CD16 | 3G8 | 157Gd | † |
| CD39 | A1 | 158Gd | † |
| TLR9 | S16013D | 159Tb | † |
| ICOS | C398.4A | 160Gd | † |
| CD303 | REA693 | 161Dy | ¶ |
| FcERIa | AER-37 | 162Dy | † |
| CCR2 | K036C2 | 163Dy | † |
| CD141 | M80 | 164Dy | † |
| FoxP3 | REA1253 | 165Ho | § |
| CD40 | 5C3 | 166Er | † |
| CD10 | HI10A | 167Er | † |
| HO-1 | HO-1-1 | 168Er | # |
| CX3CR1 | REA385 | 169Tm | ¶ |
| PD-L1 | 29E.2A3 | 170Er | † |
| CD127 | eBioRDR5 | 171Yb | § |
| NKG2D | REA797 | 172Yb | ¶ |
| TCRab | T10B9.1A-31 | 173Yb | \|\| |
| Ki-67 | 8D5 | 174Yb | ‡ |
| Tbet | 4B10 | 175Lu | † |
| CD68 | Y1/82A | 176Yb | † |
| CD57 | REA769 | 194Pt | ¶ |
| HLA-DR | L243 | 195Pt | † |
| CD103 | Ber-ACT8 | 196Pt | † |
| CD38 | HIT2 | 198Pt | † |
| CD11b/MAC-1 | ICRF44 | 209Bi | ** |

¶ Miltenyi Biotec (Charlestown, MA, USA)
||BD Biosciences (Woburn, MA, USA)
### Thermo Fisher Scientific (Waltham, MA, USA)
**Standard Biotools/Fluidigm (San Francisco, CA, USA)
