## Supplementary figures and images for "A Multigenerational “Dirty” Mouse Model for Studying Trauma-Induced Immune Dysregulation and Infection Susceptibility"

### Featured Image

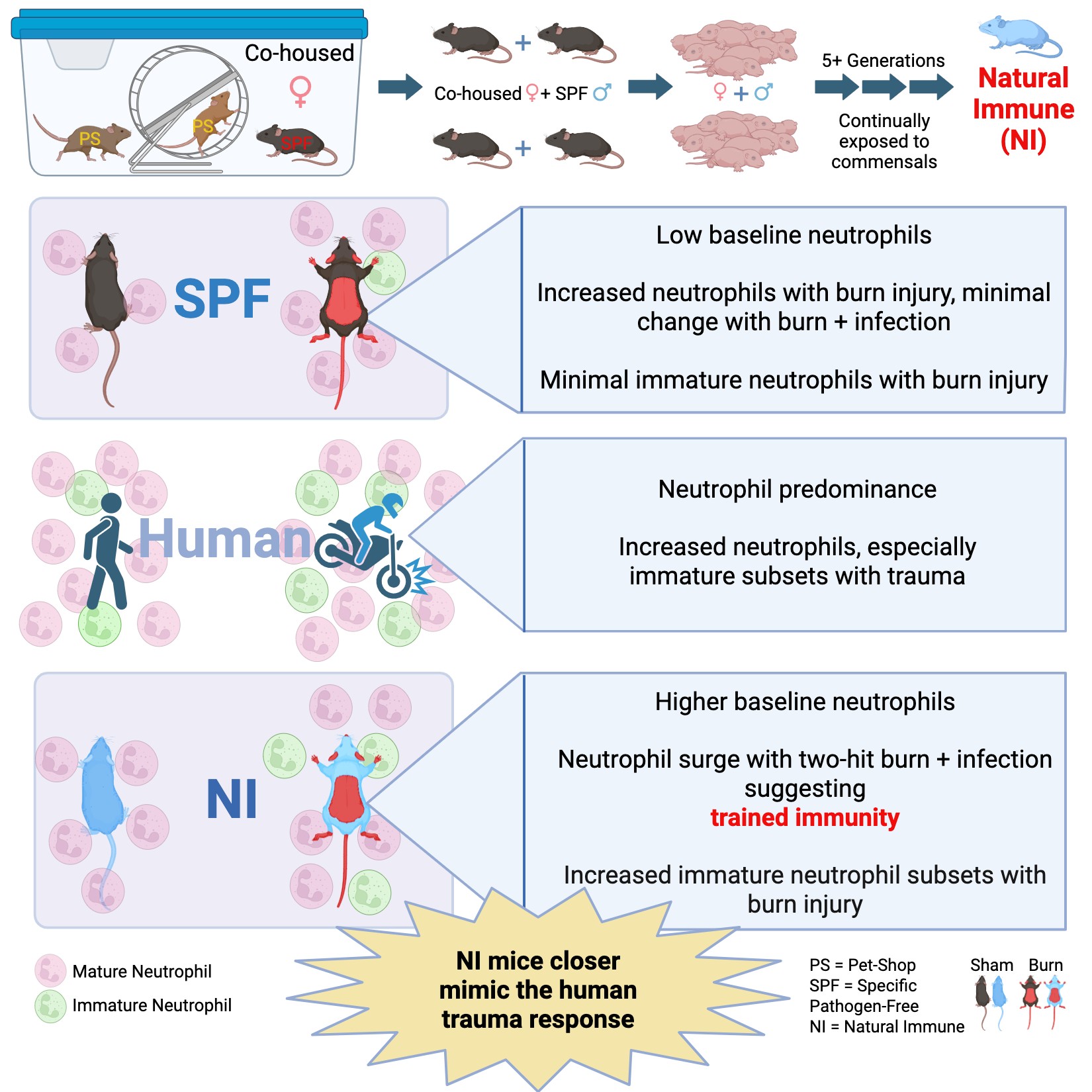
